# OSCAR: a framework to identify and quantify cells in densely packed three-dimensional biological samples

**DOI:** 10.1101/2021.06.25.449919

**Authors:** Mario Ledesma-Terrón, Diego Pérez-Dones, David G. Míguez

**Affiliations:** Depto de Física de la Materia Condensada, Universidad Autónoma de Madrid, 28049, Spain; Instituto de Fisica de la Materia Condensada IFIMAC, Universidad Autónoma de Madrid, 28049, Spain; Centro de Biología Molecular Severo Ochoa CBMSO, Universidad Autónoma de Madrid, 28049, Spain

## Abstract

We have developed an Object Segmentation, Counter and Analysis Resource (OSCAR) that is designed specifically to quantify densely packed biological samples with reduced signal-to-background ratio. OSCAR uses as input three dimensional images reconstructed from confocal 2D sections stained with dies such as nuclear marker and immunofluorescence labeling against specific antibodies to distinguish the cell types of interest. Taking advantage of a combination of arithmetic, geometric and statistical algorithms, OSCAR is able to reconstruct the objects in the 3D space bypassing segmentation errors due to the typical reduced signal to noise ration of biological tissues imaged *in toto*. When applied to the zebrafish developing retina, OSCAR is able to locate and identify the fate of each nuclei as a cycling progenitor or a terminally differentiated cell, providing a quantitative characterization of the dynamics of the developing vertebrate retina in space and time with unprecedented accuracy.

## Introduction

Cells sense, process and respond to changes in their surroundings, and the correct interpretation of these external cues is key to provide a coordinated behavior that ensures the proper function of tissues. This is of special relevance during development, when the shape, size and cellular composition of organs is established. In this context, any developing organ can be depicted as a complex multicellular dynamical system composed of many highly coordinated entities. Failure in this coordination results in the incorrect form, size and function that underlies very important developmental disorders.

The study of the intracellular and extracellular signals that regulate this complex balance is arguably one of the most recurrent topics in the field of Developmental Biology. In this direction, one of the most common approaches is based on the analysis and comparison of still microscope images, where cells can be identified by using a variety of labeling techniques. Since dynamics is an integral part in any developmental process, a more informative method involves time-lapse imaging using fluorescent proteins or dyes. Unfortunately, time-lapse imaging of developing tissues at single cell resolution is challenging due to the densely packed three-dimensional structure, specially when focusing deeper in the tissue.

Here, we present a multi-step framework (OSCAR: an Object Segmentation, Counter and Analysis Resource) designed specifically to obtain an accurate quantitative characterization of densely packed cellular tissues at single cell resolution. In brief, OSCAR uses a combination of statistical analysis, analytical geometry, nonlinear curve fitting and 3D image processing algorithms to bypass the potential errors in segmentation derived from the typical reduced image quality that results from imaging organs *in toto*.

To illustrate the capabilities of the framework, we focus on the formation of the vertebrate retina, which constitutes the most accessible part of the developing central nervous system of vertebrates, and an optimal organ to study the dynamics and the regulation of neuronal specification. The basic aspects of development of the vertebrate retina formation and the specification of its different neuronal sub-types are well characterized. In brief, a population of multi-potent retinal progenitor cells (RPCs) gives rise to several types of neurons that are produce sequentially. Retinal ganglion cells are generated first and occupy the innermost layer where they form the optic nerve. The two types of photoreceptors start to differentiate next, followed by bipolar, horizontal and amacrine cells (***Cepko et al., 1996***; ***He et al., 2012***).

Despite this apparent simplicity in its spatial and temporal organization, several key aspects of the formation of the vertebrate retina are still not understood: how proliferation and differentiation of the pool of RPCs is balanced as the retina develops?; how this balance is regulated by key regulatory signals in both space and time?; how the cell cycle dynamics of RPCs is modulated as the organ develops?; how is the interplay between cell cycle and differentiation during the formation of the retina?.

The framework OSCAR allows us to answer some of these questions by providing a highly accuarate quantitative characterization of the dynamics of growth and differentiation of the zebrafish retina. We show that that the total number of cells increases exponentially, while balancing the changes in the number of RPCs and terminally differentiated neurons. This data is then used to calculate the average mode of division and the cell cycle length, using a set analytical equations derived from a branching process formalism. The analysis shows that, during the first wave of neuronal differentiation, the balance between proliferation and differentiation change substantially. On the contrary the length of the cell cycle remains almost constant, suggesting that both mode and rate of division are independent during the first days of zebrafish retinogenesis.

The combination of a reliable and accurate tool such as OSCAR with analytical approaches to measure the mode and rate of division allows us to obtain a full characterization of the dynamics of growth and differentiation of the vertebrate developing retina with unprecedented resolution and accuracy. The straightforward application of the tool (no free parameters, no programming skills necessary) strongly simplifies the direct application of OSCAR to study the dynamics of other three dimensional tissues in quantitative detail.

## Result

### OSCAR provides an accurate quantification of in images of low signal-to-background ratio

An important problem of automated image analysis in highly dense three-dimensional biological samples is that both sharpness and signal-to-background ratio are affected by the thickness of the sample. In other words, objects closer to the surface of the tissue are sharp and well defined, but they become increasingly difficult to identify accurately as we focus deeper in the tissue. This is a direct effect of the increasing amount of scattered photons that result form the light path traveling through a thick biological sample, combined with the reduced permeability of some fluorescence dyes and antibodies into the deepest layers of a thick tissue.

In consequence, this reduced resolution highly complicates the process of automated segmentation and analysis and, even after the most extensive and careful image processing, it will inevitably result in over-segmentation (a single object identified as two or more objects) or under-segmentation events (several adjacent objects identified as a single object). Ultimately, accumulation of this errors strongly compromise the quantification of the sample at the single cell level.

A very important feature of typical 3D images reconstructed from confocal sections of three-dimensional tissues are inevitably highly asymmetric in their resolution, with the vertical axis containing much less information than the horizontal ones (i.e, the confocal plane). This is very relevant because, unfortunately, the accuracy in the quantification of 3D samples is always limited by the axis of lowest resolution. This way, even if images look sharp and nuclei can be correctly segmented in the confocal planes, the potential over-segmentation and under-segmentation errors that occur in the non-confocal planes will compromise the accuracy of automated image segmentation tools.

To illustrate this, we use images from the developing zebrafish retina fixed at two developmental stages (24 and 37 hours post fertilization, HPF) imaged *in toto* using a state of the art confocal microscope (Fig 1A-B). Nuclei (stained with ToPro3) in the XY confocal plane (left insert panels in Fig. 1A-B) appear sharp and well defined. On the other hand, nuclei in the planes perpendicular to the confocal orientation are blurry and the resolution is highly reduced (right insert panels in Fig. 1A-B), specially at later time points (Fig. 1B).

**Figure 1.**
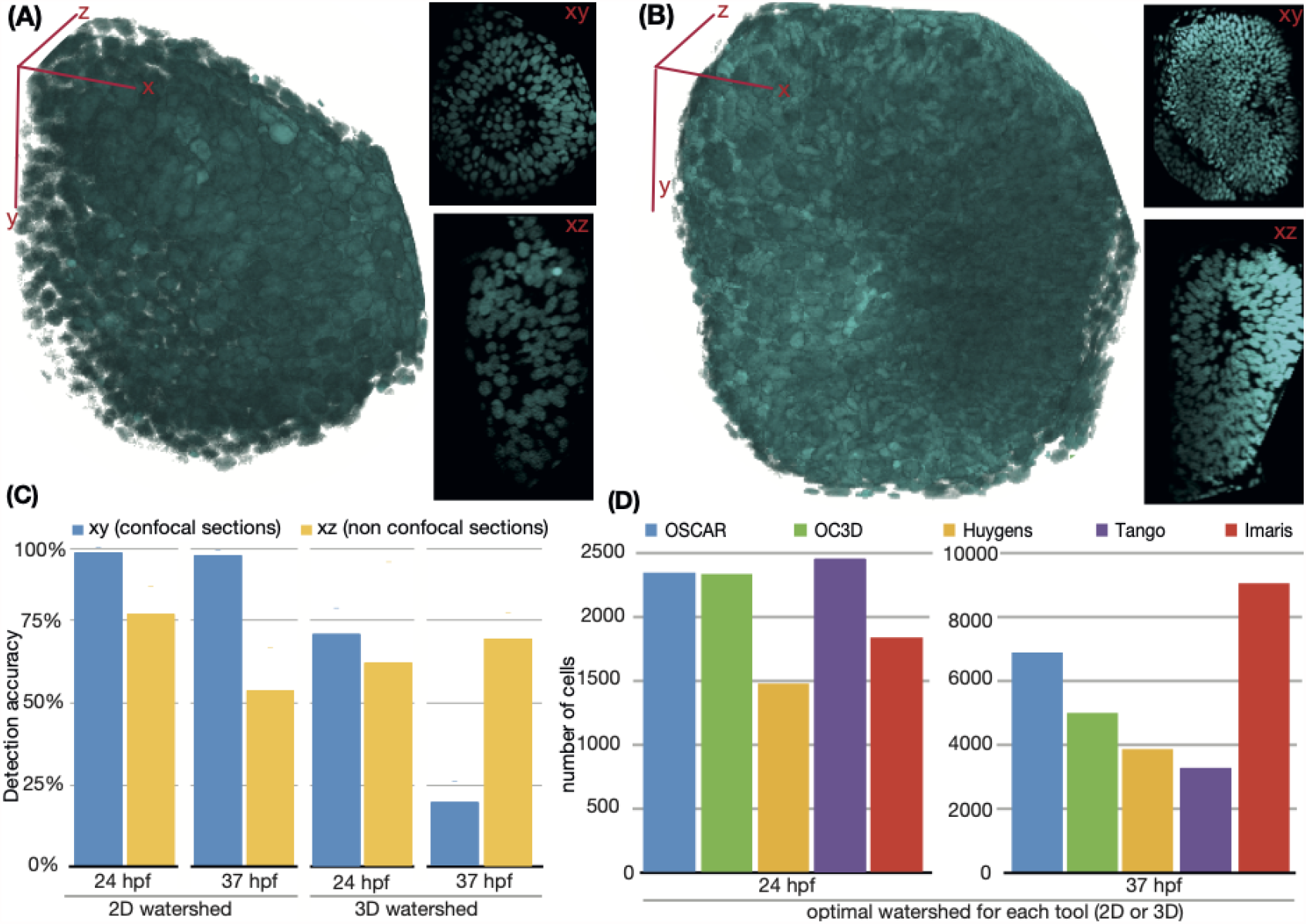
The accuracy of automated object segmentation in 3D images is limited by the axis of lowest resolution. (A-B) 3D view of reconstructions from confocal sections of the zebrafish retina stained with TopPro at (A) 24 and (B) 37 hpf. Inserts illustrate a typical confocal (above) and noncofocal (below) plane view. (C) Accuracy of object detection in 2D confocal and nonconfocal sections of the zebrafish retina using standard 2D and 3D Watershed algorithms. (D) Quantification of total number of nuclei in the 3D images above using different tools and programs.

To quantify this difference, we measured the accuracy of object detection in 2D planes in these images of the developing zebrafish retina (in terms of the number of objects detected compared to the real number of nuclei in each plane, after standard 2D watershed segmentation). Results are shown in Fig. 1C, showing that the accuracy in the confocal plane is very high at both time points (around 95%), while the automated identification of objects in the non-confocal planes (labeled as vertical) is much less precise (accuracy of 75% at 24hpf, and around 50% at 37 hpf).

With this in mind, we have designed our framework OSCAR (Object Segmentation, Counter and Analysis Resource) to overcome these situations where some axes or regions of the image have low signal-to-background ratio, compromising the quantification of the whole 3D sample. The details of the workflow of the different multi-step modules that are part of OSCAR are detailed in the Methods section.

Often, the performance and the accuracy of image segmentation and analysis tools is tested using biological images where the real number of cells is known. Fig 1D shows the total number of cells quantified with OSCAR compared to other tools commonly used in the context of 3D image quantification of biological images at cellular level. A brief description of the basics of each software tested can be found in the Methods section.

We see that, in images of lower cellular density (such as 24 HPF) OSCAR counts a very similar value number of cells that OC3D and Tango, while Huygens© and Imaris© estimate a slightly lower number. This discrepancy increases at 37 hpf (right panel), when the tissue is more crowded and potentially more segmentation errors are present. In these conditions, OSCAR identifies more cells that the other methods, with the exception of Imaris©, that counts around a 25% more cells than our framework.

Unfortunately, in thick and dense biological images such as the two examples used here, the real number of cells is unknown (is not possible to count in 3D by eye inspection with accuracy), so it is not possible to estimate the true accuracy of software solutions in these images. Often, accuracy is estimated based on “ground truth” images, where the true number of cells can be counted manually. Unfortunately, the features of biological tissues in most cases are very different from these “ground truth” samples, so the accuracy in less-than-ideal (i.e, more realistic) conditions may differ substantially than the value estimated using “ground truth” images.

To solve this problem, we generated dense artificial “ground truth” 3D tissues, composed of ellipsoidal objects (to mimic typical shape of nuclei) with size, shape and orientation obtained from gamma distributed values, with median and standard deviation defined by the user. Both location and rotation around one of the axis of the objects are chosen randomly (the details of the computer generated images are explained in the Methods section).

Next, the computer generated images are processed in different ways to decrease the signal-to-background ratio to force a level of accuracy in object detection similar to biological images, such as the zebrafish retina. Samples of these images where signal-to-noise ratio is reduced in one axis (similarly to 3D confocal reconstructions) are illustrated in Fig 2A-D. Samples where signal-to-noise ratio is decreased homogeneously are illustrated in Fig 2E, to mimic the effect of other imaging techniques such as light sheet (where loss of resolution is more homogeneous and basically due to the sample thickness).

**Figure 2.**
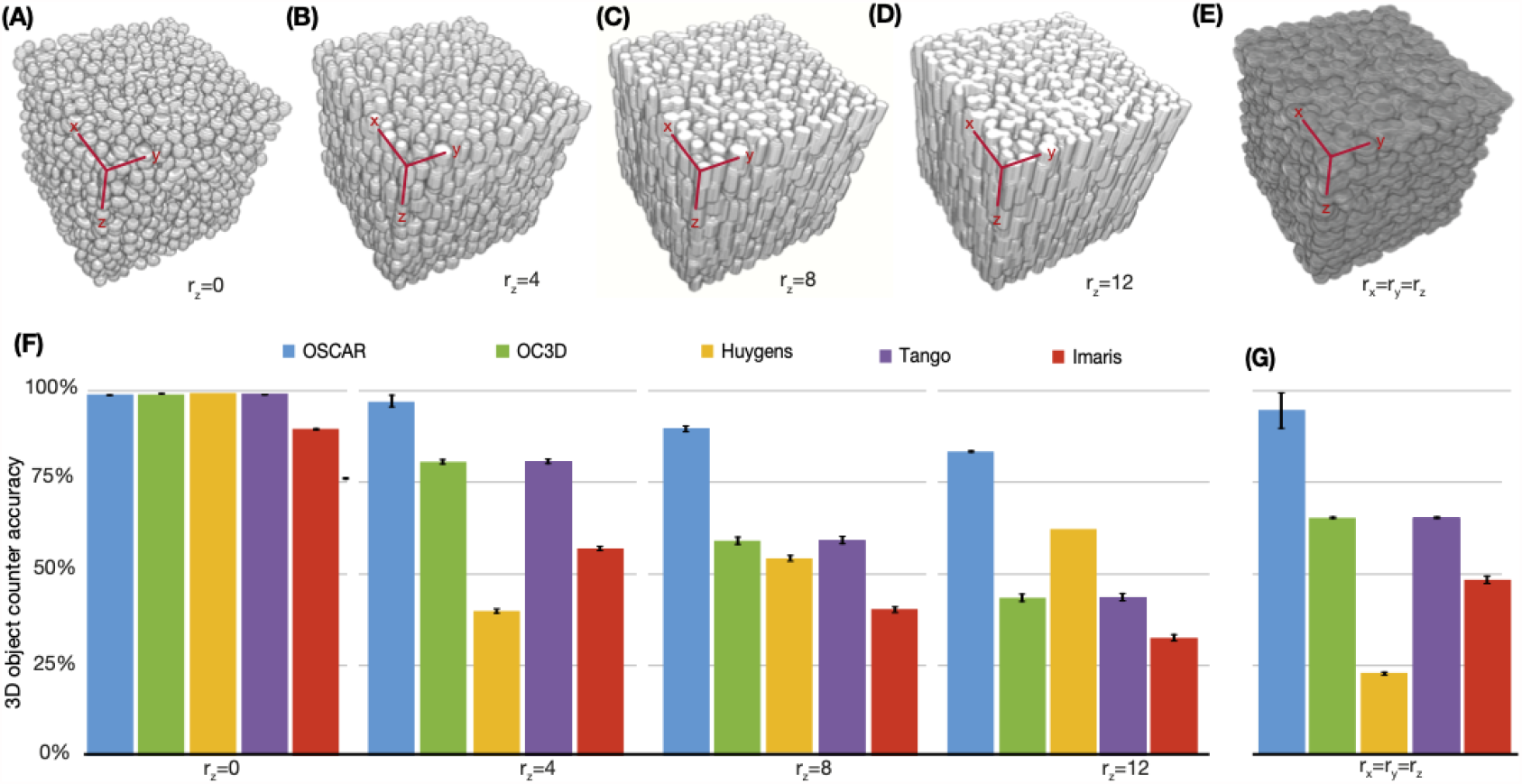
OSCAR outperforms several commonly used tools in conditions of low-signal to noise conditions. (A-D) Three-dimensional reconstruction of a set of computer generated images with 2500 cells after image processing is applied to reduce signal-to-background ratio in one axis (radius of deformation *r*_*z*_=0, 4, 8 and 12). (E) Three-dimensional reconstruction of a set of computer generated images with 2500 cells after image processing is applied to reduce signal-to-background ratio in all three axis (*r*_*x*_ = *r*_*y*_ = *r*_*z*_). (f) Accuracy in object detection in the 3d artificial stacks above for different software solutions. Error bars show the standard deviation between 3 different planes.

For the analysis, we used samples where the accuracy of segmentation in *z* (see Supp Fig 1) is similar to the accuracy when segmenting the nonconfocal planes of the zebrafish retina (Fig 1C (Fig 2C, yellow bars) after automated segmentation using 2D Watershed.

Results of the total number of objects detected in the different conditions by OSCAR and other image analysis tools are summarized in Figs. 2F-G. All tools provide an accurate quantification of the total number of objects in conditions of high signal-to-background ratio (radio of deformation, *r*_*z*_=0). As the signal-to-background ratio is decreased to levels where the segmentation accuracy is similar to the biological 3D images of the zebrafish retina (*r*_*z*_=8, *r*_*z*_=12), OSCAR is able to detect objects with an accuracy above 80%, while other tools report a number of objects that is around 2X above or below the real count (50% accuracy, one out of two objects is). In addition, OSCAR also outperforms (90% accuracy) the other solutions when the signal-to-background is reduced homogeneously in all directions (Fig. 2G).

In conclusion, our data shows that OSCAR outperforms other tools commonly used in the field of biological image analysis in situations, in conditions of highly dense cellular environments and low-signal-to-background ratio. The difference in performance occurs both when signal-to-noise is reduced in one axis only (similar to confocal reconstructions) or homogeneously (similar to light-sheet images). This better performance is possible because OSCAR is designed to take advantage of the 3D information for each object to bypass potential errors of segmentation, providing a much improved capability to identify and analyze biological samples where accurate automated segmentation is not possible, such as *in toto* 3D tissue samples and/or *in vivo* time lapse movies.

### OSCAR accurately localizes objects in 3D space

Another very important feature of image analysis tools is their ability not only to count, but to identify the correct location of the objects in the sample. A good localization is key to quantify key features of the sample, such as local density, cell tracking between frames of a taime-lapse movie, and even identification of cell identity based on data from other channels of the image. To visually inspect the accuracy of OSCAR and compare it with other tools, we plot the objects detected (orange spheres) on top of the 3D images of the zebrafish retina (blue) at 24 and 37 hpf (in Fig. 3A). At first inspection, the objects detected by OSCAR, OC3D and Tango presents less clustering of their output objects and with a higher degree of similitude with the real images of the retina.

**Figure 3.**
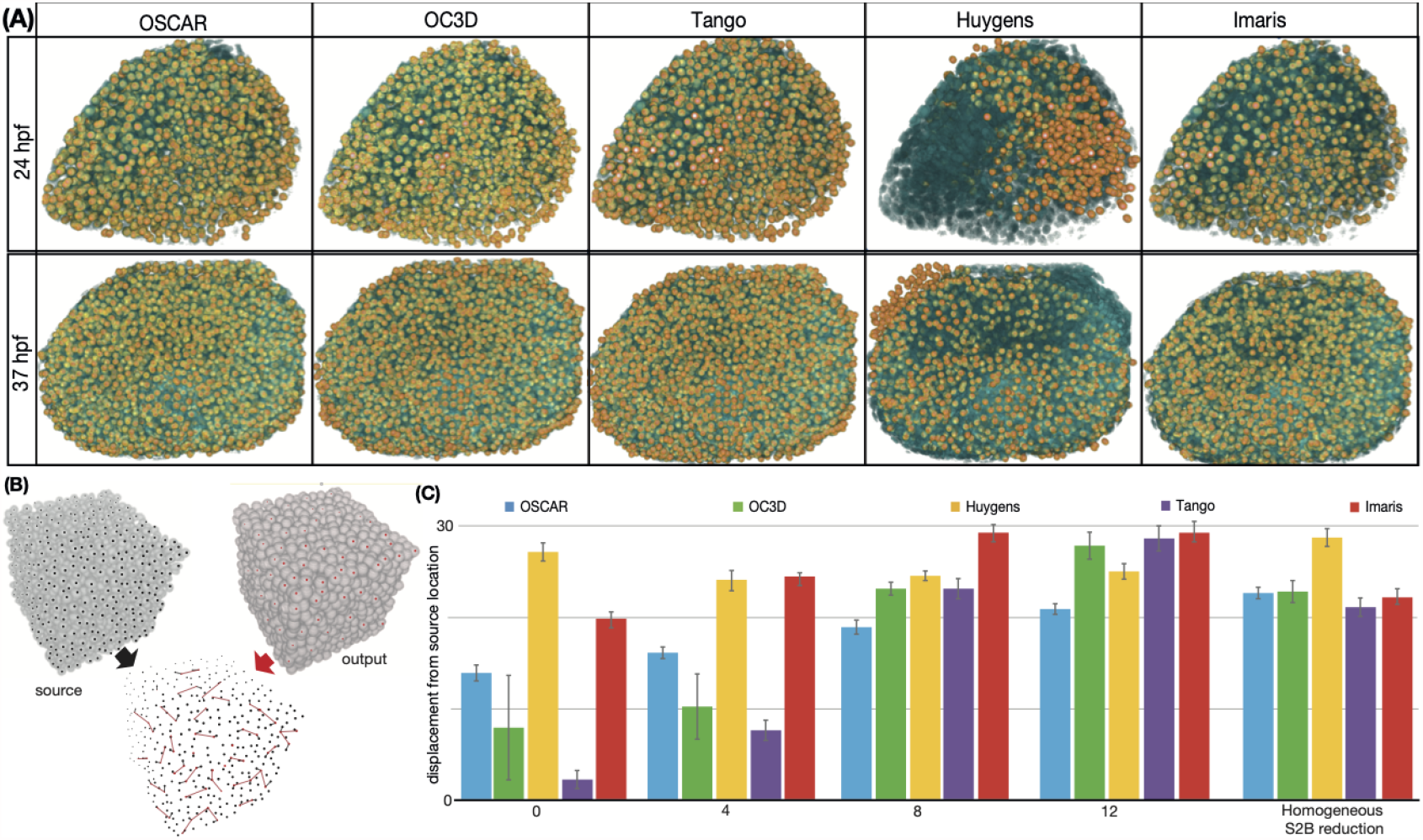
The OSCAR framework applied to biological samples localizes individual cells with high accuracy. (A) Overlapping of the biological images of zebrafish retinas a different time points (blue) and the output of OSCAR and other tools tested (orange). (B) Scheme of the algorithm used to quantify localization accuracy of the different tools. (C) Quantification of accuracy in the spatial localization of objects by the different tools over computer generated images using a Friedmann-Rafsky test. Error bars represent the standard deviation, resulting from multiple independent runs of the algorithm.

A more quantitative characterization of the accuracy in the detection of objects in the 3D space can be performed by using again computer generated images, since the location of all objects on the sample is known. To do that, we develop an algorithm based on the Friedmann-Rafsky test that compares the difference in location of two sets of points in a 3D space (illustrated in Fig. 3B, details of the algorithm are explained in the Methods section). In brief, the algorithm establishes links between objects of the output image and their closest neighbours in the input image. Next, the sum of the euclidean distance of all the links (red lines in Fig. 3B) is used to compute the deviation between the location of the objects in the input and output images.

To correct for the difference in the number of total objects predicted by the different methods, we only use a subset of objects from the output in the algorithm. The process is repeated multiple times for different subsets of centroids, and the mean values are plotted in Fig. 3C (smaller values indicate less distance between location of objects of input and output, and therefore, more accurate spatial detection).

The plot shows that the localization accuracy for all tools tested is reduced as the signal-to-background ratio decreases, as expected. Although Tango and OC3D are highly accurate when images are sharp, OSCAR outperforms the other software solutions in conditions of more noisy images (i.e., more segmentation errors). When image resolution is reduced homogeneously (Fig 2E), almost all tools show similar accuracy than OSCAR, despite the fact that they are able to detect only around half of all objects in this image (Fig. 2G).

The higher accuracy in detecting the correct number of objects, and the higher accuracy in locating them in the 3D space, opens up the very useful possibility to identify each cell in crowded tissue as a different cellular sub-type based on potential information from other channels of the image. This feature is illustrated in Fig. 4, where we use an image that combines nuclear staining (ToPro3) in zebrafish retinas with Phospho-Histone H3 immunostaining, a marker for cells in Meta phase of the cell cycle), in green (Fig. 4A). The green channel of the image is processed (see Methods) to identify nuclei as positive or negative for phosho-histone-3 immunostaining.

**Figure 4.**
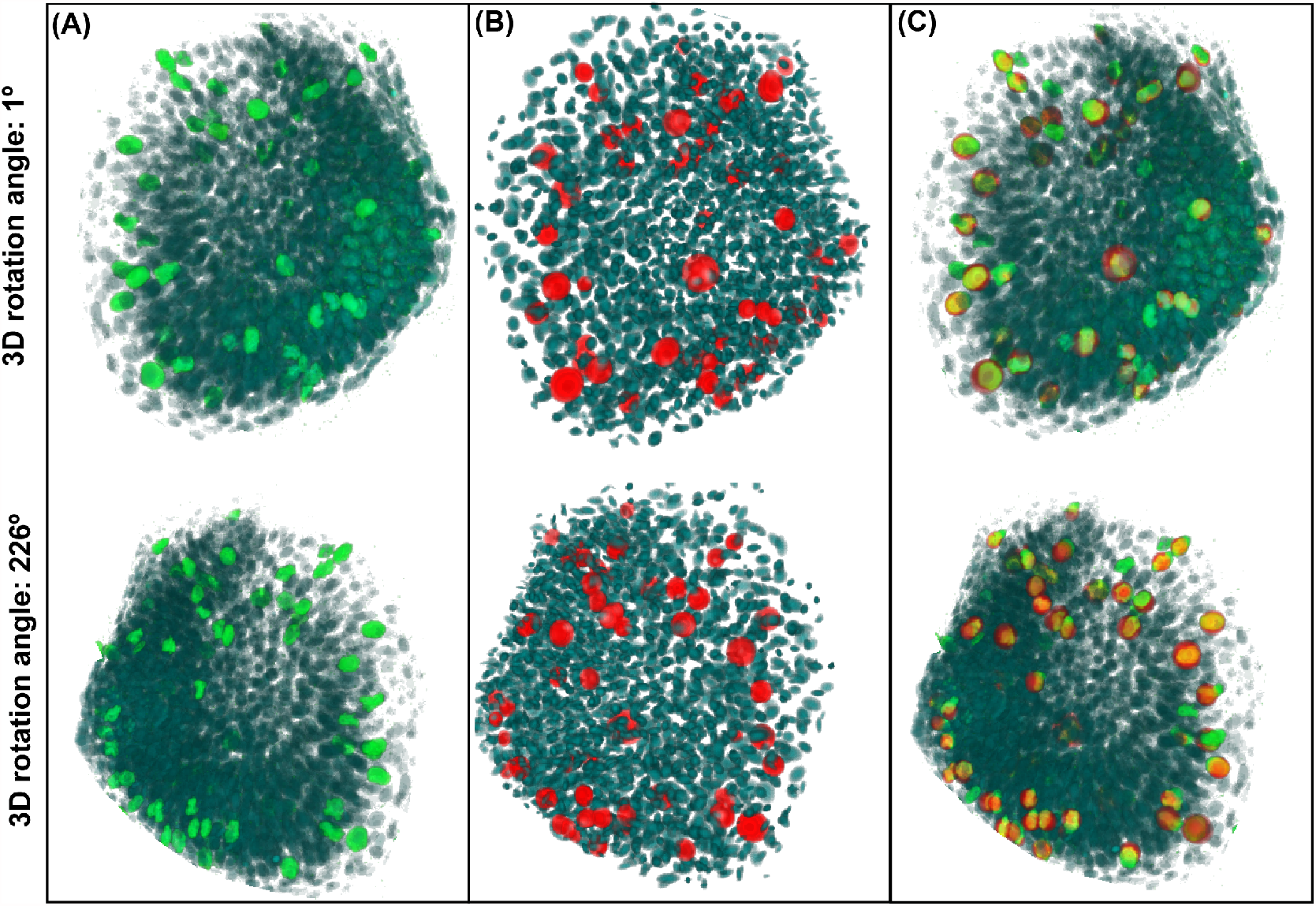
Location of phosho-histone 3 positive cells in the developing zebrafish retina. (A) 3D image reconstruction of the zebrafish retina at 24 and 37 hpf stained with nuclear marker (blue) andphosho-histone3 immunostaining (green). (B) Digital representation of the same image from the output of OSCAR where each nuclei detected is represented as a 3D ellipsoid (blue). Location of cells that are detected as phospho-histone3 positive is plotted in red. (C) Superposition of the detected cells in M-phase on top of the input image (A).

Output of OSCAR is shown in Fig. 4B, where the objects detected as positive for Phospho-Histone H3 immunostaining are labeled in red. Fig 4C, overlaps only the red objects of the output on top of the input image (the one shown in Fig. 4A). The high overlap between green (real cells im M-phase) and red (cells detected by OSCAR in M-Phase) illustrates the high accuracy in establishing celular identity in dense 3D images.

In conclusion, the capability of identifying all objects in a sample with higher accuracy, coupled to the ability to correctly locate these objects in the 3D space, opens up new possibilities for accurate automated image analysis of tissues composed of different cell subtypes with features that can be distinguished by specific labeling. This feature of OSCAR will be used in the next sections to quantify the dynamics of development of the zebrafish retina.

### Quantification of the dynamics of the developing zebrafish retina

In this section, we demonstrate the capabilities of the framework applied to the characterization of developing zebrafish retina in four dimensions (three dimensions + time). A preliminary attempt to characterize the dynamics of the formation of the zebrafish retina in space and time was performed in Ref. (***Li et al., 2000***), where authors estimated the number of total cells in the zebrafish retina using a very clever method based on reconstruction of slices and the average size of the nuclei. Another excellent approach to study the vertebrate retina in 3D is presented in Refs. (***Pan et al., 2013***; ***Almeida et al., 2014***), where transgenic zebrafish lines are engineered using the Brainbow technology to study the global dynamics of differentiation and the interaction between different cell types. Unfortunately, authors did not quantify cell numbers, nor the details of the differentiation dynamics. In addition, Icha et al. (***Icha et al., 2016***) use 3D information to characterize key aspects of the dynamics of retinal ganglion cells after differentiation. Recently, (***Matejčić et al., 2018***) used three-dimensional images and theoretical analysis to determine that changes in cell height maintain the shape of the developing zebrafish retina throughout development. In this contribution, authors measure the number of cells and the number of differentiated neurons at different developmental times using FACS.

To quantify the dynamics of the developing zebrafish retina, we proceed as follows (a scheme of the process is illustrated in Supp. Fig. 2): embryos of the atho5:Egfp zebrafish line are allowed to develop in standard conditions (see Methods) and are fixed at different developmental time points; whole retinas are then processed, cleared, mounted, stained with nuclear marker and immunofluorescence against EGFP and SOX2 to label progenitors and terminally differentiated neurons; whole embryos are imaged *in toto* using a confocal microscope; finally, the images (at least three independent replicates per time point) are processed and analyzed using OSCAR (details of the each part of the protocol can be found in the Methods section).

Examples of nuclear staining (ToPro3) in sections and 3D reconstructions of retinas at three different hours post fertilization (hpf) are shown in Fig. 5A. These images are used as input of OSCAR to identify all cells in the tissue. The corresponding sections and 3D reconstructions are provided as output of OSCAR are shown below each image, where cells represented as ellipses with the shape, volume and location in the 3D space provided by OSCAR. Direct comparison of the two images shows a good agreement between the real retina and the digital reconstruction.

**Figure 5.**
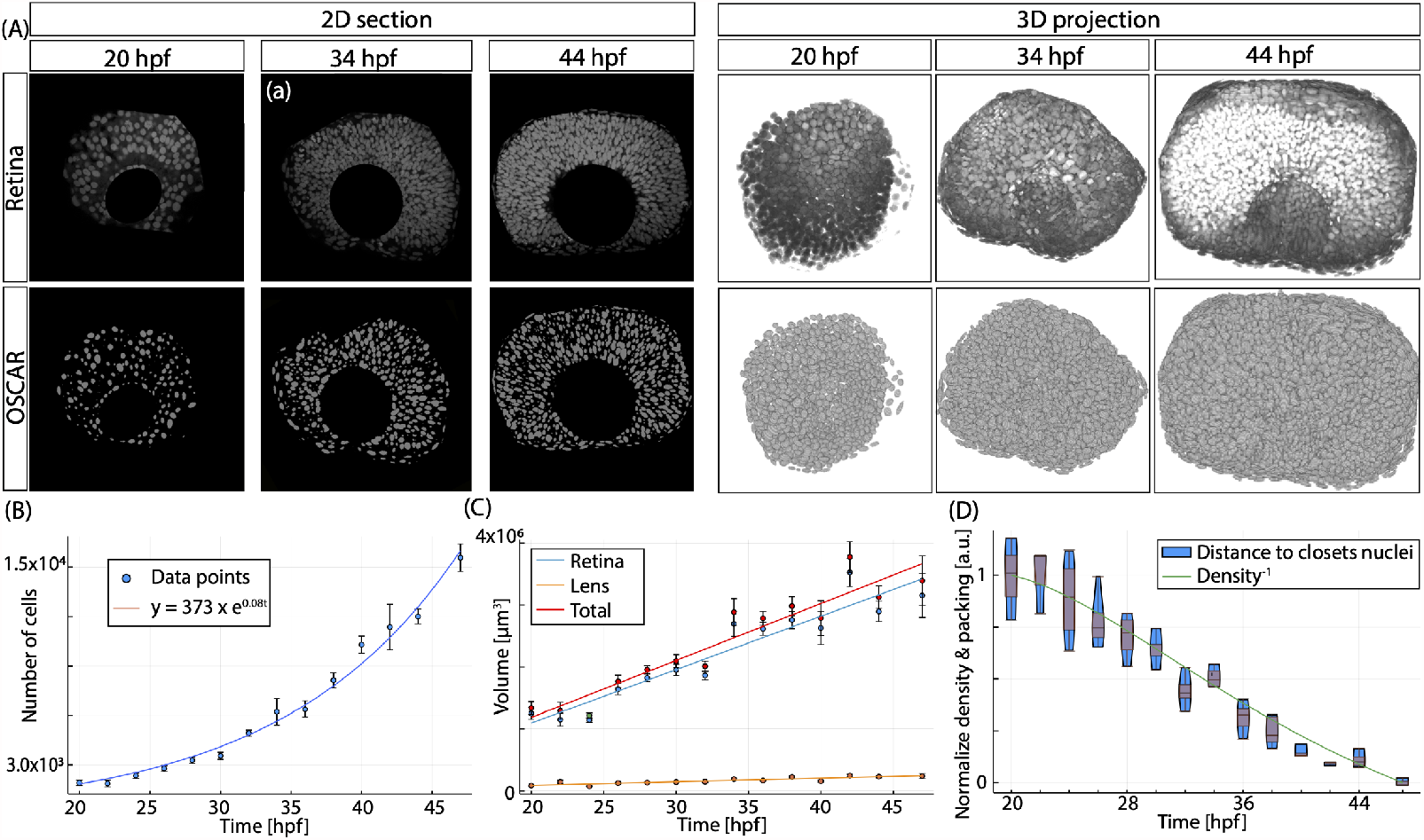
Quantification of the growth of the wild type zebrafish retina using OSCAR. (A) Representative confocal sections and 3D reconstructions of developing zebrafish retinas at different time points stained with nuclear marker ToPro. Below each image the corresponding digital representation is plotted, generated as output of OSCAR (cells are drawn as ellipsoids with dimensions and location for each cell provided by OSCAR). (B) Plot of the total number of cells at different developmental stages (at least three independent repeats per time point). Data is fitted to an exponential curve (line). Error bars represent the standard error of the mean value between repeats. (C) Estimation of volume occupied by the retina and lens as the tissue develops. Error bars represent the standard error of the mean value between repeats. (D) Box-plot of the average distance between 5-closest neighbors. Line represents the change in density measured as the fitting for number of cells divided by the fitting for the volume of the retina.

The total number of cells quantified using OSCAR is shown in Fig. 5B, where data seems to follow an exponential evolution (blue line). This is inconsistent with previous findings where recent studies with less temporal resolution report two distinct regimes with linear growth dynamics (***Matejčić et al., 2018***).

Interestingly, although the number of cells increases exponentially, the volume of the retina increases more linearly (Fig. 5C). This is inconsistent with recent findings where authors observed a more exponential growth in the volume (***Matejčić et al., 2018***). We hypothesize that this discrepancy may be due to their reduced temporal resolution (6 hours between time points). On the other hand, the doubling time in volume reported (around 12.5 h) is consistent with our values at the initial points in our analysis (the volume doubles in 12 hours between 20 hpf and 32 hpf).

The difference between the linear increase in volume or the exponential increase in cell numbers suggesting that the tissue has to become more dense as it develops. This change in density can be estimated from the spatial location of each cell in the tissue provided as output of OSCAR. To do that, we compute the average distance of each cell in the tissue to its five closest neighbors The results are shown as a boxplot in Fig. 5D, where we see that the change in density is more prominent between 28 to 36 hpf. Interestingly, this is consistent with the calculation of the density obtained by dividing the linear fitting of the temporal evolution of the volume of the retina (Fig. 5C, blue line) by the exponential fitting of the total number of cells (Fig. 5B, blue line). The inverse of the value of the density is shown also as a blue line on top of the boxplot. The good agreement between both quantities suggests that both the number of cells and the location provided by OSCAR are consistent with each other.

### Identification of progenitors and differentiated cells allows to quantify the dynamics of the cell cycle and differentiation

Next, by tanking advantage of the accuracy in object detection in the 3D space of OSCAR, we can identify each cell in the tissue depending on its fate as progenitor or terminally differentiated cell. To do that, we use the information on the other color channels of the confocal images, that correspond to the immunostaining against SOX2 and EGFP (details of this process are explained in the Methods section). Fig. 6A illustrates a representative section of a zebrafish retina at 34 hpf, where we mark with arrows nuclei where SOX2 staining is strongly decrease in cells that are expressing GFP, while few other cells show high GFP expression and also SOX2 expression (marked by yellow arrows).

**Figure 6.**
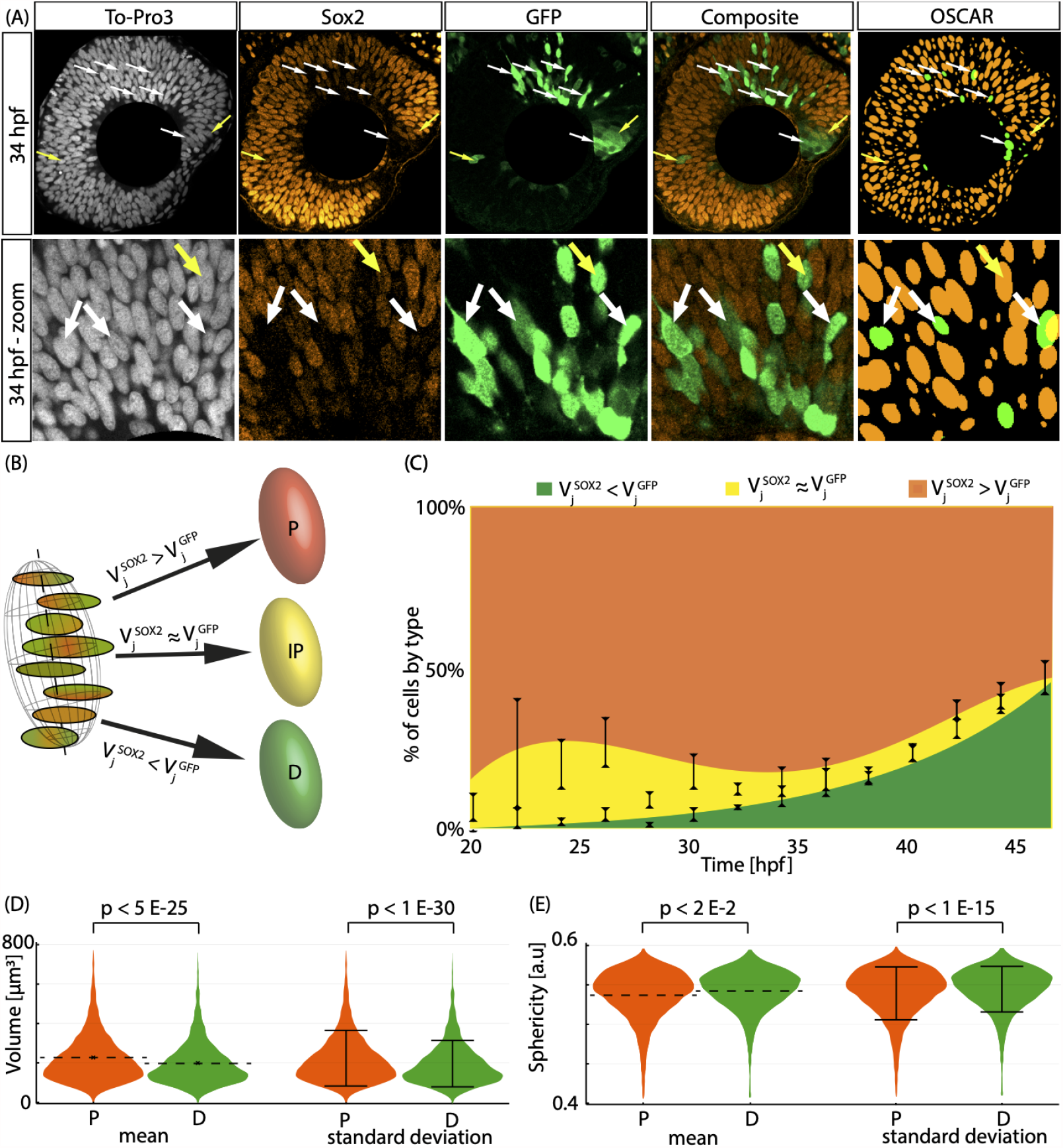
The identity of the cells in the zebrafish retina can be established based on the combination between immunostaining intensities. (A) Different channels of an image of a representative section of a developing retina at 34 hpf. A zoom of each image is shown below, to illustrate cells positive and negative for each staining. (B) Scheme of the decision process to establish identity of the 3D objects detected by OSCAR.(C) percentage of cells in the retina where SOX2 staining intensity is measured as higher (red), lower (green) or similar (yellow) that GFP immunostaining. (D-E) Quantification of differences in size and sphericity between nuclei of progenitors and differentiated cells identified by our algorithm.

Next, we establish the identity of each object detected by OSCAR as positive or negative for each particular staining by comparing the amount of pixels inside the 3D-object that are above certain threshold value for each immunostaining (details of the algorithm that establishes the identity of the objects are explained in the Methods section). In brief, a cell is identified as progenitor or differentiated if is the number of pixels above a threshold for SOX2 is significantly higher than for GFP, an vice-versa (Fig. 6B). The quantification for each type at the different time points is shown in Fig Fig. 6C, where we can see that, initially a large percentage of cells fulfill the condition 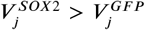, and therefore can be safely labeled as Progenitors. The percentage of cells in the tissue that fulfill 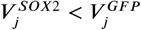, and that can be identified as terminally differentiated, increases as time progresses.

On the other hand, there is a significant amount of cells where staining intensity is similar for both channels, (yellow cells) and therefore, their identity cannot be established simply by comparison of the two channels. In these conditions, the identity has to be established based on additional estimations or previous studies.

To establish the identity of the yellow cells, we performed immunostaining against PCNA (a marker of cells that are actively proliferating) and Phosho-histone 3 (which labels cells that are undergoing m-phase), at different developmental times. We used OSCAR to quantify the cells that are positive for each of these staining as well as GPF. Results are shown in Supp Fig 3 and Supp Fig 4 for PCNA and PH3, respectively. We can see that a small percentage of cells that are GFP+ are also PCNA+, suggesting that some cells starting to express Ath5 are still actively cycling. The same is true when quantifying PH3, we see a small but non-negliglibe amount of cells that are labeled as positive for GFP+ that are in M-phase.

This is consistent with previous studies where authors monitored closely the activation of the Ath5 promoter in this zebrafish line, reporting the existence of ath5:GFP progenitors that later become photoreceptors, amacrines, or horizontal cells (***Poggi et al., 2005***). In addition, recent studies show that neurogenic progenitors that arise from asymmetric divisions are Ath5 positive, suggesting that these are reminiscent of the intermediate progenitors found in the mammalian neocortex(***Nerli et al., 2020***). In our analysis, we observe that the percentage of these intermediate progenitor cells is high at initial time points, and then decreases, suggesting that, in average, the percentage of asymmetric divisions is higher at early stages than a later stages of the first wave of differentiation of the developing zebrafish retina.

Finally, we can use the features measured by OSCAR to investigate the morphological differences between cells identified as progenitors and differentiated. Estimation of the differences in volume is plotted in Fig 6D, where we observe that the nuclei of differentiated cells are smaller in volume than in progenitors, with high statistical significance. In addition the standard deviation is also difference, suggesting less variability in the volume of the nuclei of differentiated cells, again with high statistical significance. We can also monitor the sphericity of the nuclei (how similar to an sphere are the nuclei detected). Results shown in Fig Fig 6E show a similar mean, with small but statistically significant differences, while the standard deviation in differentiated cells is again reduced, compared to progenitors. Overall, our data shows that nuclei of differentiated cells are smaller and more consistent in size and shape than nuclei of progenitor cells.

### The cell cycle and the mode of division vary during the first wave of differentiation

In the present section, we will use the output of OSCAR to investigate the changes in cell cycle and differentiation dynamics. To do that, multiple independent tissues (at least three per developmental stage) are processed and stained with To-Pro3, SOX2 and GFP immunofluorescence, as explained in the Methods section. Examples for three developmental stages are illustrated in Fig. 7A for a representative section and a whole mount 3D-reconstruction. The digital corresponding images resulted form the output of OSCAR are shown below, showing good agreement with the images of the retina, both in sections and 3d-images comparison.

**Figure 7.**
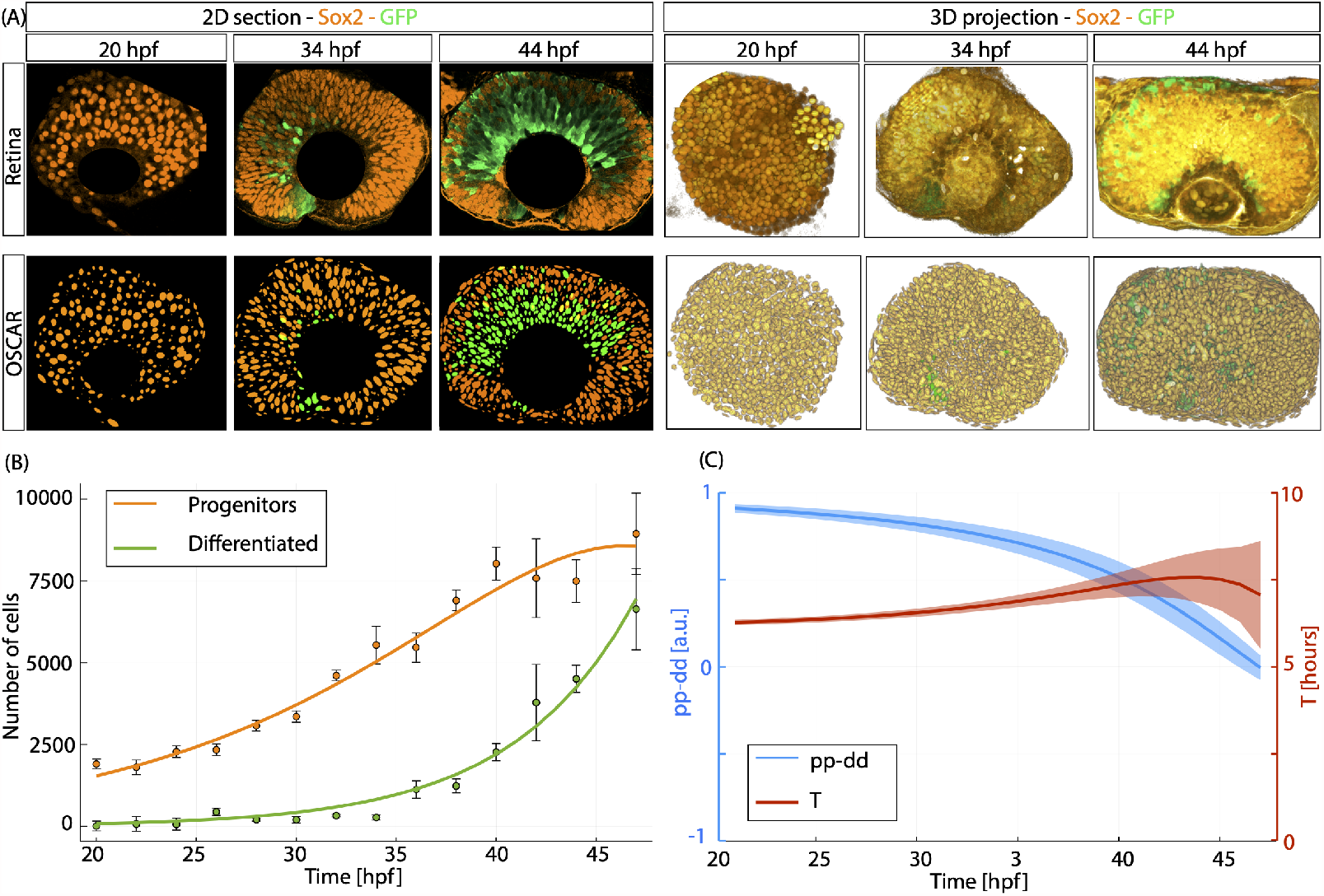
Quantification of the dynamics of proliferation and differentiation of the wild type zebrafish retina using OSCAR. (A) Representative sections and 3D-view of tissues stained with SOX2 (orange) and GFP (green) at different developmental stages. Below each image, we plot the corresponding digital representation to illustrate the output of OSCAR. (B) Quantification of the number of progenitors (orange dots) and differentiated cells (green dots) overtime. Data from progenitors is fitted with a two parameter double-exponential (green line) *P* (*t*) = *p*_1_ · *exp*(0.1 · *t* + *p*_2_ · *exp*(0.1 · *t*)), *p*_1_= 224; *p*_2_= -0.01. Differentiated cells (green line) are fitted with a two-parameter exponential, *D*(*t*) = *d*_1_ * *exp*.(*d*_2_ · *t*), *d*_1_=3.10; *d*_2_= 0.164. Error bars illustrate the standard error of the mean value for each time point. (C) Average cell cycle length (red, right vertical axis) and average mode of division (blue, left vertical axis) obtained from equations 6-7, when using as input the fitted curves *P* (*t*) and *D*(*t*). Ribbons illustrate the confidence interval, calculated from based on the standard error of the mean value for progenitors and differentiated cells.

To extract the value of the average cell cycle and the mode of division, we take advantage of a set of analytical equations developed by our group and published previously in Refs. (***Míguez, 2015***; ***Ledesma-Terrón et al., 2020***). In brief, this equations based on a branching process Markow chain formalism, provide values of the average cell cycle length and mode of division in populations of cells where proliferation and terminal differentiation is being taking place simultaneously. The equations and the details of the branching process tool are explained in the Methods.

The branching process formalism requires as input absolute values of the total number of progenitors and differentiated cells in the retina. Therefore, based on our own data and the previous literature indicate, intermediate progenitor cells (both positive for SOX2 and Ath5) are still cycling progenitors, and therefore are considered as such for the purpose of this study.

The total number of progenitors (red dots) and differentiated cells (green dots) in the retina at different developmental stages obtained as output of OSCAR are plotted in Fig. 7B. Interestingly, the dynamics of differentiated cells fits well to an exponential growth mode (green line), while the progenitors (red line) exhibit an initial exponential increases until 35 hpf, when the growth dynamics is slowly reducing concomitant with with the increase in the rate of production of the terminally differentiated cells.

Data from differentiated and progenitors are fitted by two parameter exponential and double exponential, respectively. Predicted values from this fitting are then used as input of equations 6-7, together with the value of the growth fraction and the apoptosis rate. The growth fraction *gamma* is estimated based on immunostaining against PCNA (see Supp Fig 3), which is commonly used to mark cells that are actively cycling. The time evolution of *gamma* is assumed as a linear change between the two data points measured. For the rate of cell death during this stages, it has been shown previously (***Matejčić et al., 2018***; ***Dzafic et al., 2015*)** that apoptosis is negligible between 20 hpf and 48 hpf. Therefore, the value of *emptyset*_*p*_ is set to zero for the present analysis.

Output of the equations is plotted in Fig 7C. We can see that changes in the mode of division (blue line, left vertical axis) are significant, starting with very little differentiation that gradually increases to a point that reaches a balance between the production of new progenitors and the loss in the population due to differentiation. This region where *pp* − *dd* ≈ 0 is consistent with the values observed in Fig 7C for the final time points, where the amount of progenitors remains almost constant.

Values for the average cell cycle in the progenitors are shown in red (right vertical axis), suggesting that the cell cycle remains almost constant with a slight increase overtime from 6.5 hours to 7.5 hours, but this change may be due to high variability in the values of progenitors and differentiated cells at later time points, that result in broader intervals of confidence.

## Discussion and Conclusions

## Methods and Materials

### Animals

Experiments are performed in Tg (ath5:GFP) zebrafish line, engineered to express eGFP as a reporter for Ath5 expression, a very well characterized biomarker of early RGCs (***Poggi et al., 2005***). The Ath5:eGFP zebrafish line was sustained according to the standard procedures and protocols (***Westerfield, 1994*)**. All experimental protocols were performed in accordance with the guidelines of the European Communities Directive (2012/63/EU) and Spanish legislation (Real Decreto 53/2013).

The embryos are incubated in E3 1X fish medium (5 mM NaCl, 0.17 mM KCl, 0.33 mM CaCl2, 0.33 mM MgSO4) supplemented with Methylene Blue (Sigma) at 28°C. Pigmentation of cells in the Retinal Pigmented Epithelium that surrounding the Neural Retina is blocked by adding 0.003 % phenotiurea (PTU;Sigma) at 22 hpf directly to the E3 1X medium (***Westerfield, 1994*)**. Culture media is replaced every 24 hours.

Experiments proceed as follows. Embryos are initially collected at one cell stage in petri dishes filled with water where animals are maintained. Next, embryos are washes with E3 1X fish medium several times for cleaning. Next, embryos are separated into sets of 20 per each p100 petri dish and placed in an incubator at 28° C humidified to minimize evaporation of culture media.

Fertilization time is controlled by separating females and males the previous day (***Westerfield, 1994*)**. Embryos at similar developmental stage developmental are classified based on visual inspection. Next, embryos are collected at different developmental stages (***Kimmel et al., 1995***) and fixed in a solution of Formalin at 10% (Sigma, HT501128) overnight at 4°C.

After fixation, embryos are washed several times with PBS and the chorions are manually removed. Finally, embryos are dehydrated using serial dilutions of methanol at 25%, 50%, 75% and 100% in PBS.

### Immunohistochemistry

Immunoflourescence staining is performed then *in toto* using a protocol adapted from Ref. (***Cer-*** *veny et al*., *2010*). Briefly, embryos are re-hydrated using serial dilutions of 75%, 50%, 25% and 0% methanol in PBS.

All solutions used in this protocol are performed using a solution of PBS 1X and 0.6% triton (PBT 0.6%) as the solvent, to increase permeability of antibodies and dyes. Next, to further maximize antibodies permeability inside all regions of the threedimensional tissue, samples are treatment with proteinase K (1 mg/ml). The duration of the treatment is adjusted for each particular developmental stage (from 15 at 22 hpf to 45 hours at 48 hpf).

Next, embryos are treated with a short pulse of Formaline at 10% (Sigma, HT501128) to preserve the integrity of the tissue from mechanical manipulation during the immunofluorescence process. Next, after Formaline is washed several times from the tubes,

Next, blocking is performed using a solution of the Fetal Bovine Serum 10% (FBS) diluted in PBT 0.6% for 1 hour at room temperature or overnight at 40C. Next, tissues are exposed to the primary antibodies diluted in a solution of FBS 2% in PBT 0.6% in a weighing balance overnight at 4°C.

After primary antibody exposition, several washes of the sample are performed (five minutes for each one).

Next, tissues are exposed to the secondary antibodies and other fluorescent molecules as nucleic acid or membrane counterstain, diluted in a solution of FBS 2% in PBT 0.6% for 1 hour at room temperature. After second antibody exposition, several washes of the sample are performed (five minutes for each one).

The primary antibodies used in the immunofluorescence of zebrafish embryos are: chicken anti-GFP (1:1000, Abcam ab137827), rabbit anti-Sox2 (1:1000, GeneTex GTX124477), mouse anti-PCNA (1:100; Invitrogen, MA5-11358) and rabbit anti-PH3 (1:250; Merck-Millipore, 06-570). And for the secondary antibodies: goat anti-Chicken-488 (1:500; ThermoFisher, A-11039), goat anti-mouse-55 (1:500; ThermoFisher, A-21157) and donkey anti-rabbit-555 (1:500; ThermoFisher A-31573). DNA (and, subsequently, the nuclei) is stained with interkalant nuclear agent 647 nm fluorophore To-Pro3 (1:500; ThermoFisher, T-3605) for all samples.

### Image Acquisition

Light refraction, scattering and dispersion when traveling through thick biologically dense samples quickly reduces the quality of the image. Therefore, the minimization of this effect is key to obtain high quality images with high signal-to-noise ratio in all confocal planes of the image.

To minimized the amount of biological tissue that the light needs to travel before reaching the sample, whole eyes are manually dissected from the whole embryos before mounting. Next, eyes are mounted in in-house made chambers (to minimize changes in the shape of the eye) with RapiClear 1.49 © (SunJin Lab). A coverslide is carefully is placed on the sample to be visualized with confocal microscopy.

Images of confocal planes of the sample (1024×1024 pixel resolution) are taken in a Leica SM800 confocal microscope, using 1 *µ*m of pinhole. An overlapping region between confocal slices of 0.2 *µ*m is established to ensure a correct reconstruction of the tissue in three dimensions.

### Image Processing of retina confocal sections

Prior to analysis, images are enhanced by sequential application of filters and processing to sharpen and enhance the contrast. The set of filters used depends strongly on the image, and is designed to facilitate the 2D segmentation in the 2d planes that then are used by OSCAR to reconstruct the objects in the 3d space. This is performed using FIJI (***Schindelin et al., 2012***). For the images of the developing zebrafish retina in this contribution, the sequence of processing algorithms and filters used was the following.

1. We define the size of the Kernel Radius (*KR*), i.e., a parameters that defines the square-shaped local area where several of the filters and image processing algorithms used below are applied locally over the whole image image. The *KR* was fixed as 2.5.
2. Local increment of differences between the foreground and background intensities applying a local kernel window of size 8x*KR*x*KR* that was moving across the image, removing all pixels with a value less than the median of each local kernel window.
3. The previous filter separates adjacent 2d-objects, but also generates holes and breaks the integrity of objects. To solve it, the following sequence of filters is applied to remove noise and increase the definition of the boundaries of each 2d-object: Gaussian Blur filter, Maximum Filter, Median filter and Unsharp Mask filter.
4. A binary mask was generated by applying a threshold (median) intensity as cutoff value (all pixels where the intensity is below the threshold are set to zero).
5. Euclidean Distance Map (EDT) was performed in the binary image to generate the primordium points (seeds) for all detected objects. Seeds are then used by a flood fill algorithm to find boundaries of each object (***Kang et al., 2010***).
6. Objects smaller than *KR*\**KR* are discarded from further analysis.

### OSCAR

This section explains the sequence of processes implemented in the framework to reconstruct the 3D objects from the data of the 2D confocal planes.

First, each 2d-object in the confocal planes is fitted to a parametric 2D function of interest. Since OSCAR is used here to fit nuclei from biological images with a mainly 3D-ellipsoidal shape, sections in XY planes are therefore best fitted by to ellipses. The parameters of the ellipse associated to each object, such as area, centroid location, orientation and excentricity, are used to compute the values used to establish the interactions between objects in the planes. By relying on the parameters from the fitting, OSCAR is able to perform calculations and statistical analysis to establish the interactions between objects of different planes orders in a fraction of time and computational cost compared to using the full shape of the object.

To initiate the plane-by-plane sequential reconstruction of the 3d-Objects, the framework starts by identifying the first ellipse of a given 3d-Object. Establishing the correct identity of these “seeds” is key in the process of reconstruction and in the final number of objects identified, and therefore, it has to be implemented carefully.

To do that, the framework determines if a given generic ellipse *i* of the plane *z* is identified as the initial section of a 3d-object based on the interactions of the ellipse *i* with the ellipses in the planes above. This characterization can be performed in different ways, depending on the characteristics of the image, in terms of nuclei density and signal-to-background ratio. Some of the potential possibilities implemented are illustrated in Fig. 8. First, the identification as “seed” can be established based on the distance to sections in plane *z* − 1 (Fig. 8A). In this approach, a given ellipse *i* is identified as a “seed” if it is not chosen among the k-closest ellipses of any ellipse in the plane above. A less restrictive approach is based on the quantification of the intersection between fitted ellipses in neighboring planes. This intersection is calculated computationally by computing the set of points inside ellipse *j* in plane z-1 that fall inside ellipse *i* in plane *z* with foci in 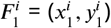 and 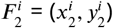, and major axis *a*_*i*_. Therefore, a point (*x, y*) inside of ellipse *j* is also inside of ellipse *i* when the following inequality is fulfilled.

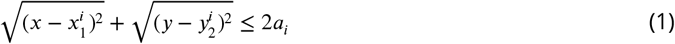

the intersection *U*_*i,j*_ is calculated as the sum of all the points (*x, y*) in ellipse *j* that fulfill this condition.

**Figure 8.**
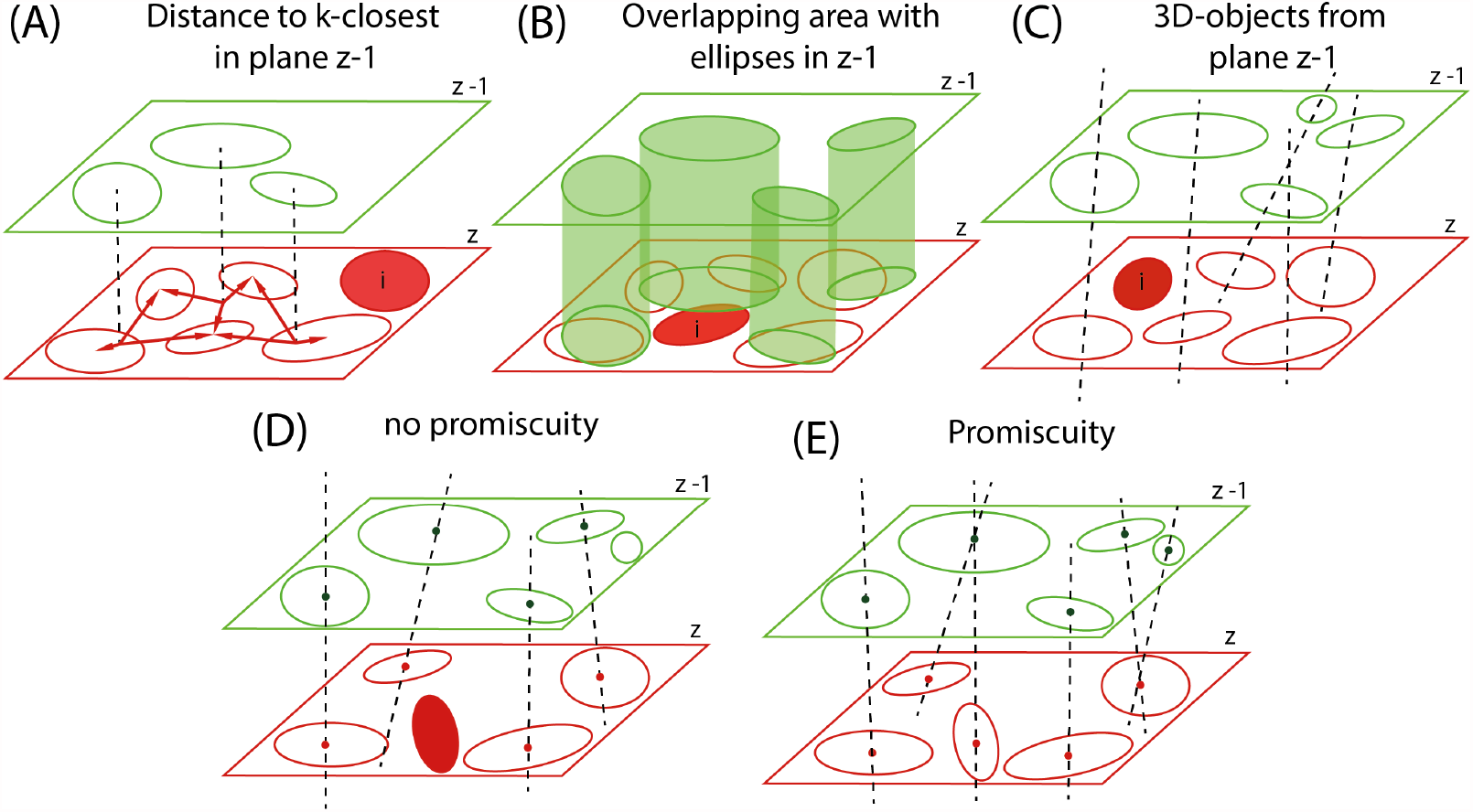
Identification of ellipses as parts of 3D objects. (A-C) identification of ellipses as initial sections of 3D-objects (marked in red). (A) The ellipse *i* has not been chosen among the k-closest ellipses of any ellipse in the plane z-i. (B) The ellipse *i* does not overlap above a certain threshold value with any ellipse of the plane z-1. (C) The ellipse *i* is not part of any previously formed 3D-object. (D) Each fitted ellipse can be only part of a single 3D-object. (E) Each fitted ellipse can be part of multiple 3D-objects.)

**Figure 9.**
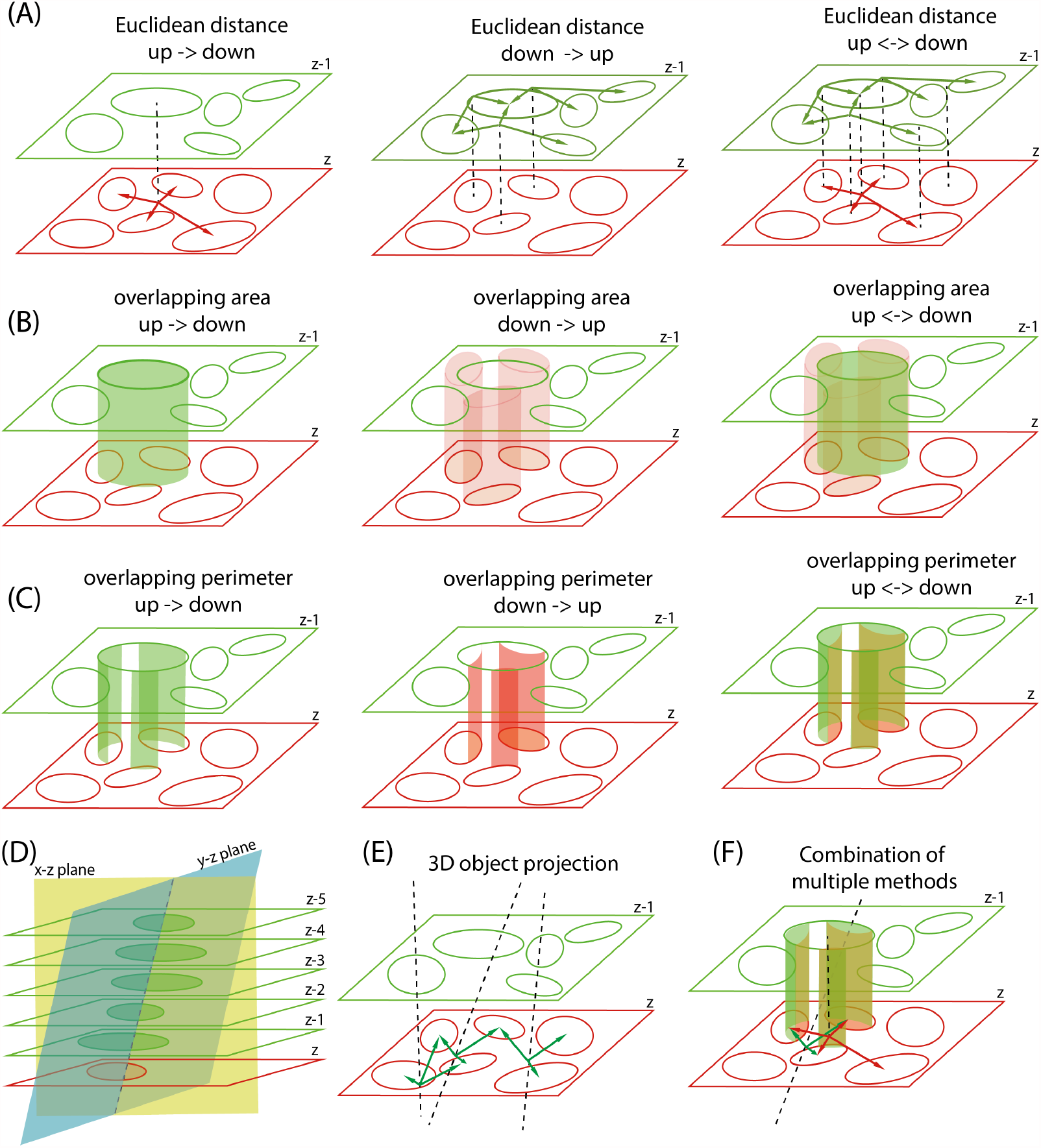
3D object reconstruction techniques used by OSCAR. (A) Object reconstruction based on Euclidean distance between centroids of ellipses in consecutive planes. This way, for a given ellipse *i* in plane z-1, calculation can be computed from: up-to-down (closest ellipses in z); down-to-up (ellipses in z that choose the ellipse *i* as their closest); bidirectional (choose ellipse that are closest to *i* and also choose *i* and one of their closest) (B) Object reconstruction based on area overlapping: up-to-down (area from *i* into ellipses of plane z); down-to-up (area from ellipses in z into area of ellipse *i*; bidirectional (which corrects for bias due to the fact that ellipses have different sizes). (C) Object reconstruction based on perimeter overlapping. This is equivalent to the previous case, but computationally less costly. (D-E) Euclidean distance to 3D object projection in plane z. The planes *xz* and *yz* are generated based on location of centroids of ellipses in object. Ellipses in z are chosen based on the distance to the intersection between *yz* and *yz* planes. This approach account for the fact that objects may not be aligned in *z*. (F) Object reconstruction based on combination of several methods concatenated. For instance, choose the k-closest based on Euclidean distance (A), filter the selection by bidirectional perimeter overlapping (C); finally filter by Euclidean distance to 3D object projection (E). This combination has been used to quantify the images in the present manuscript.

**Figure 10.**
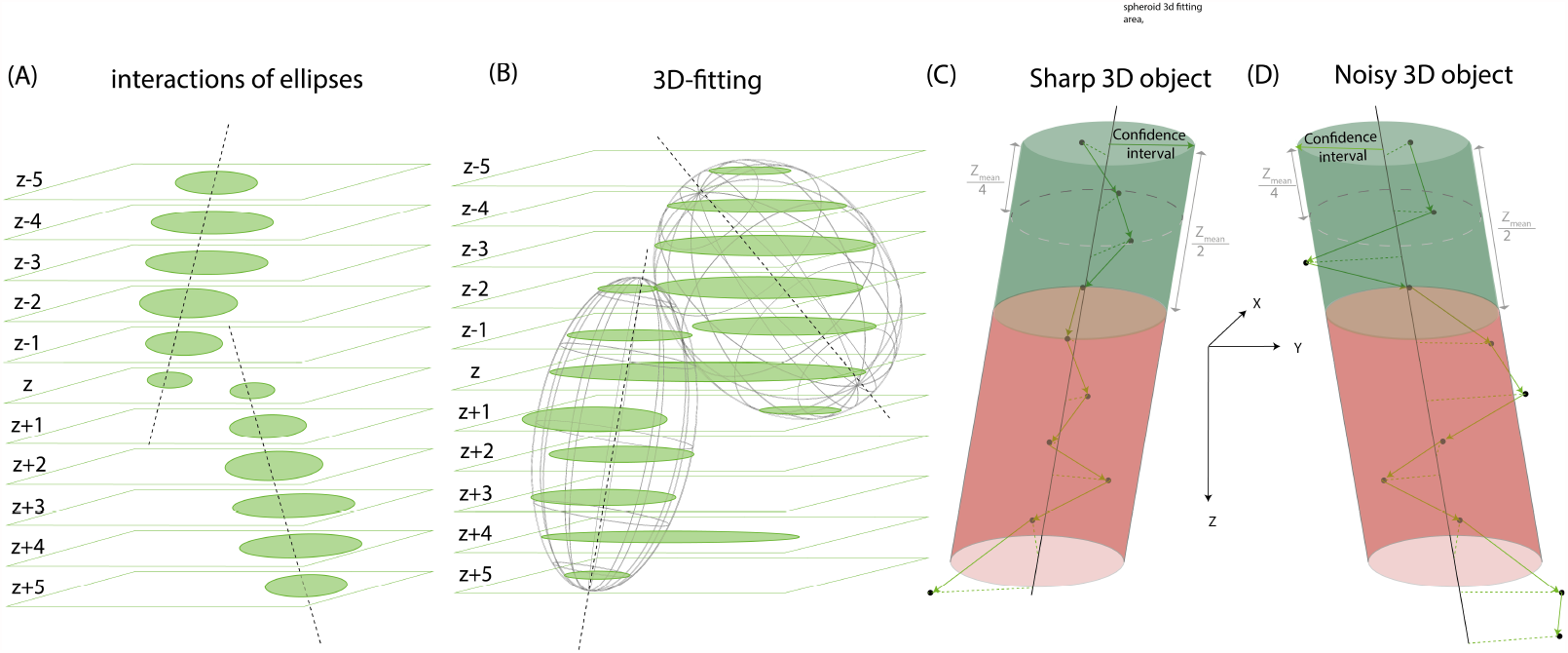
Different processes to establish end points of 3D object. (A) In sharp images or not dense tissues, the last ellipse of a 3D-object that does not interact with an ellipse in the following plane is considered the end point of the Object. (B) When some segmentation errors are present, a good approach is to use the first half of the 3D-object to fit a 3d-parametric curve and estimate the location of the last section of each object. (C) Identification of end points based on confidence intervals. The last centroid inside the confidence interval established is the end point. (D) For very noisy objects, the final point is established when more centroids after the average half side are outside the confidence interval than before the average half size of the 3D-objects.

**Figure 11.**
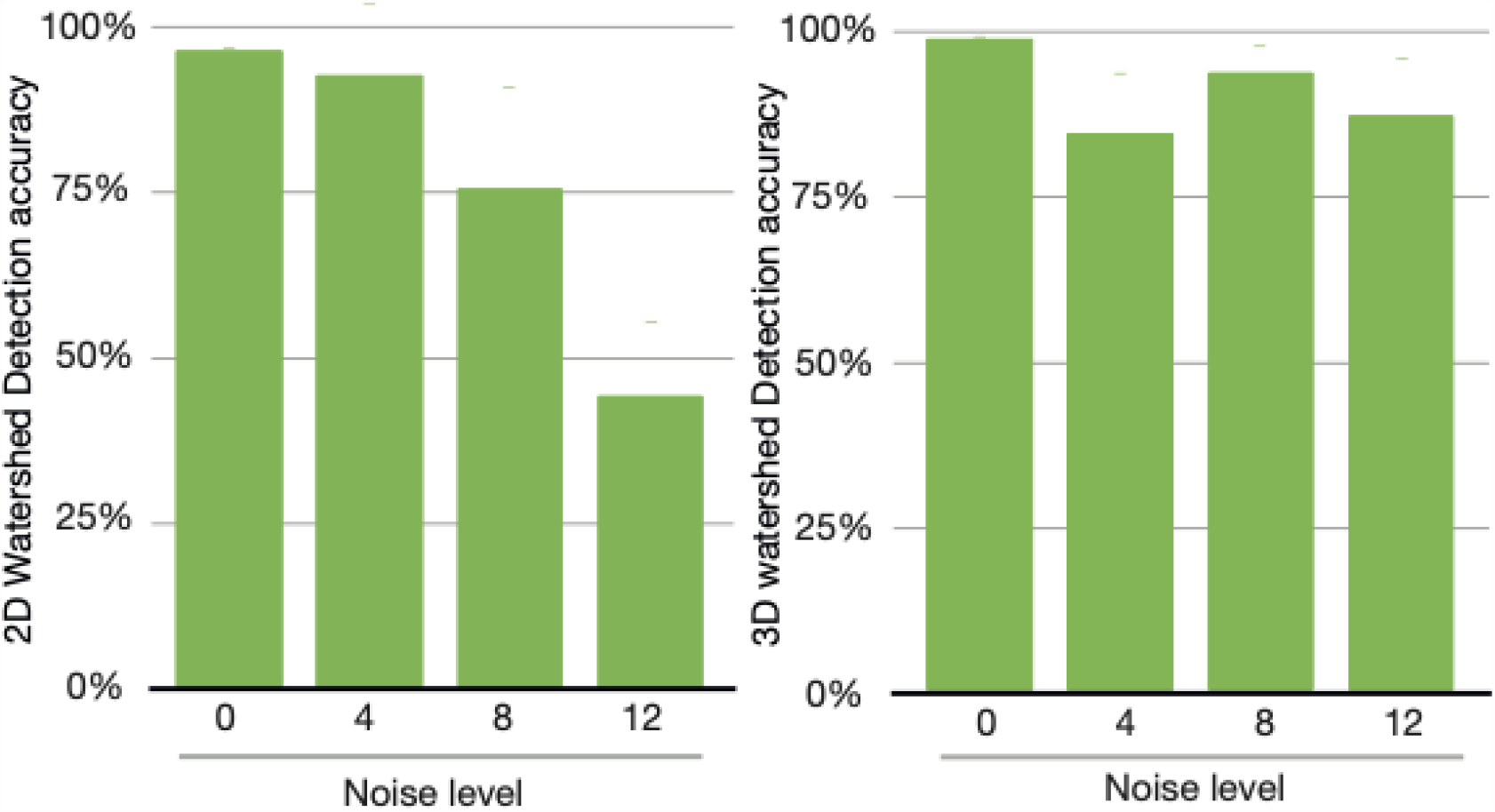
Supplementary Figure 1: Detection accuracy in artificial images of different noise levels after 2D or 3D watershed segmentation.. Segmentation is performed using FIJI stock algorithms for 2D or 3D watershed.

**Figure 12.**
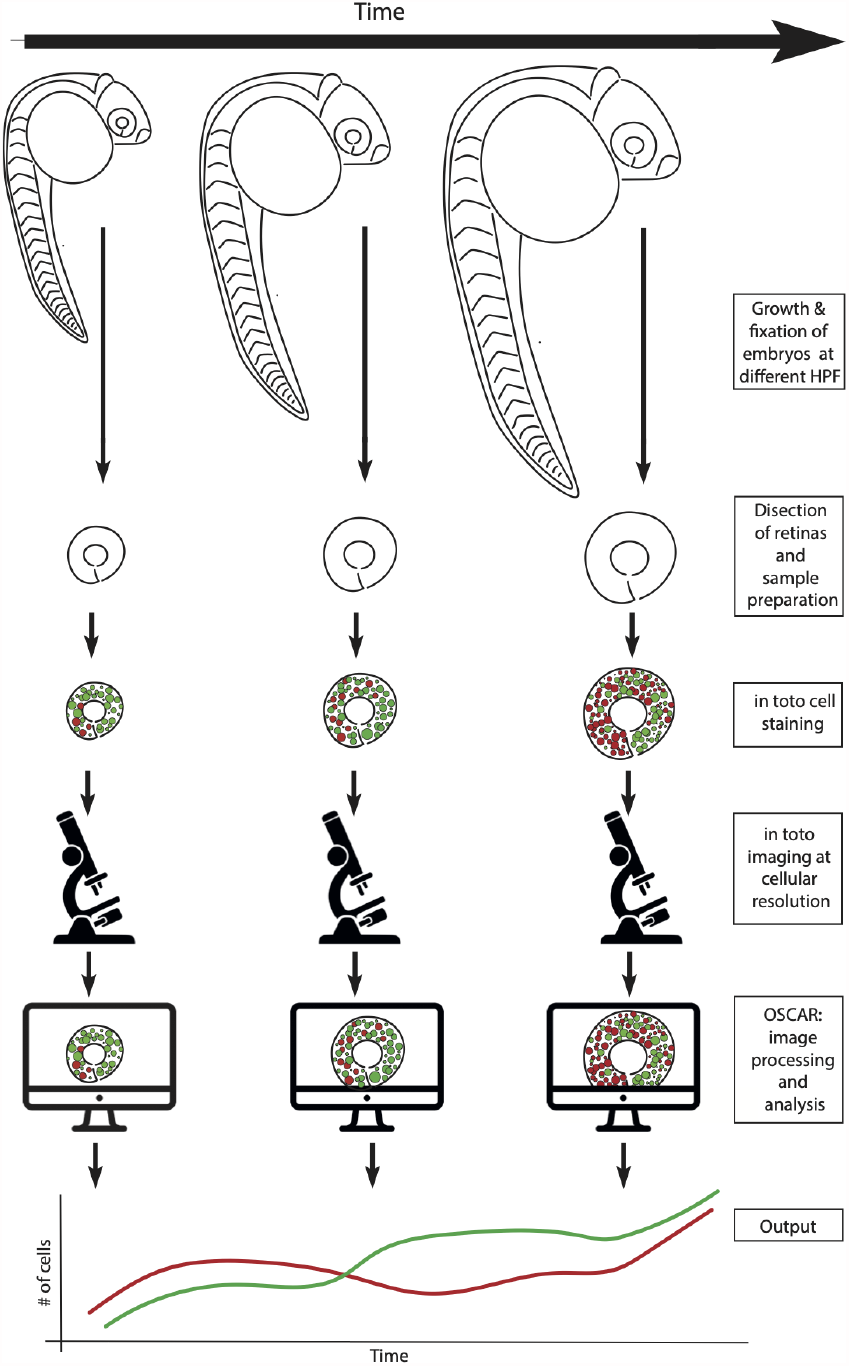
Scheme of the experimental protocol. Embryos are allowed to develop in normal conditions and fixed at different time points. Retinas are then dissected, and immunofluorescence is performed *in toto* using the combination of antibodies and/or fluorescent staining desired for each experiment. Next, retinas are imaged *in toto* in a confocal microscope with 30% overlap between planes to obtain an optimal 3D reconstruction of the tissue. Finally, the multiple independent repeats for each condition and time-point are processed and quantified using OSCAR. Output of OSCAR is used to monitor key features of the development of the Zebrafish retina, such as density, number of cells and proliferation and differentiation dynamics.

**Figure 13.**
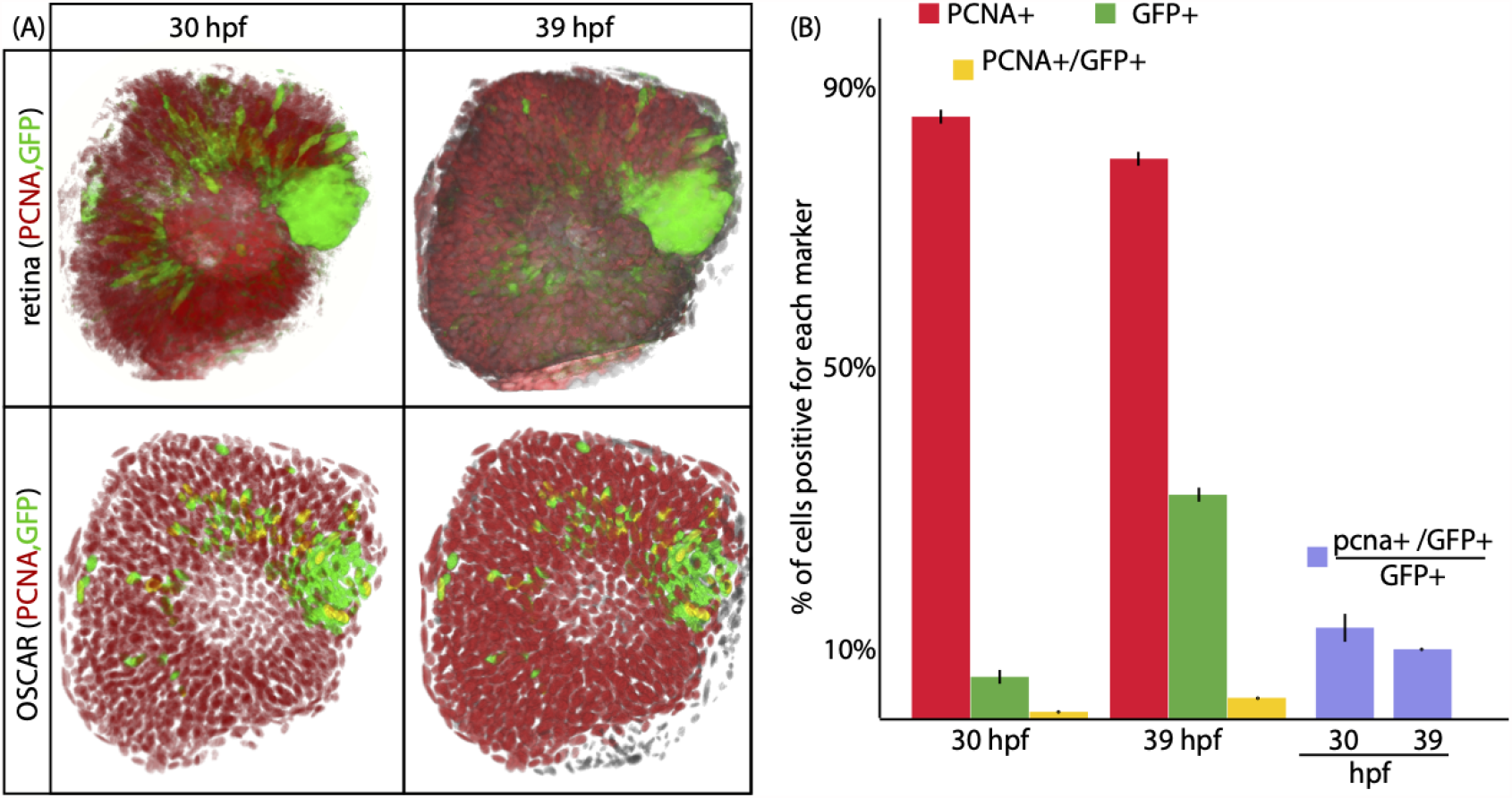
Supplementary Figure 3: *in toto* quantification of PCNA in the zebrafish retina. (A) 3D-view of the whole zebrafish retina at 30 and 39 hpf stained with PCNA and GFP. Below each image, the digital version of the image above represented as output of OSCAR. (B) Quantification of the number of cells that are PCNA+, GFP+ and double positive for PCNA and GFP.

**Figure 14.**
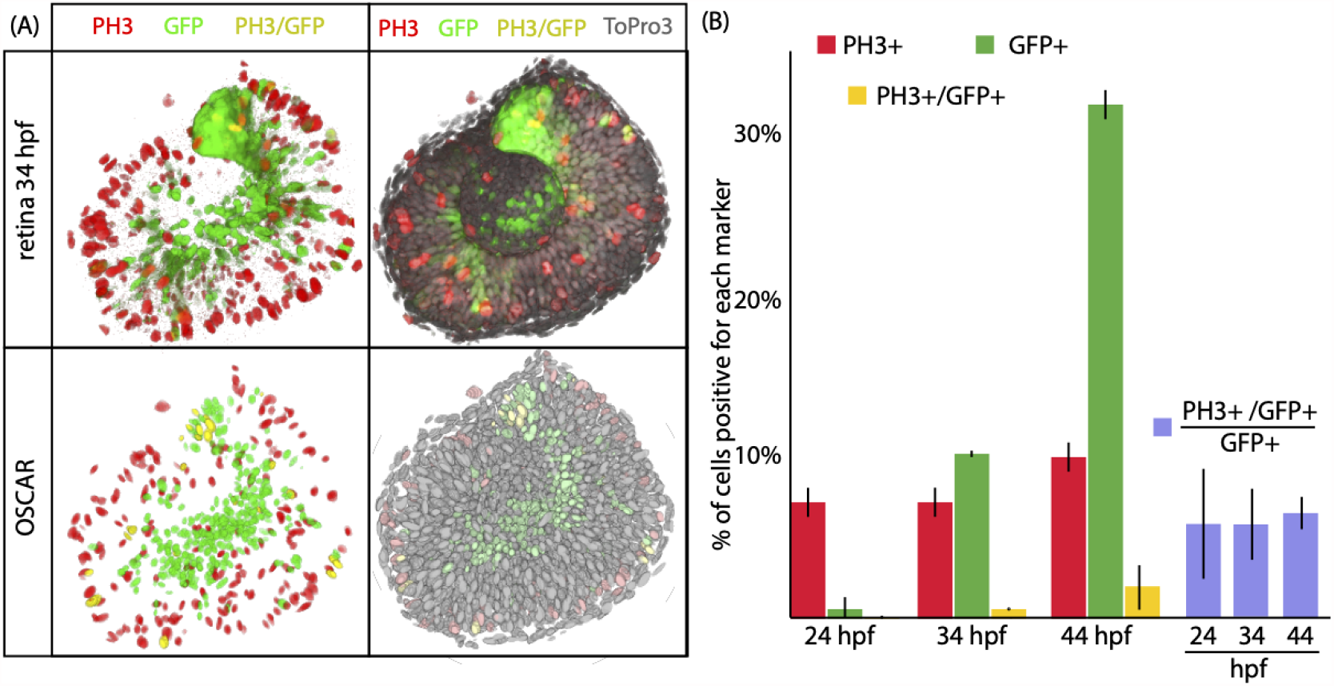
Supplementary Figure 4: *in toto* quantification of Phospho Histone 3 in the zebrafish retina. (A) 3D-view of the whole zebrafish retina at 34 hpf stained with Phosho histone 3, GFP and To-Pro3. Below each image, the digital version of the image above represented as output of OSCAR. (B) Quantification of the number of cells that are PH3+, GFP+ and double positive for PH3 and GFP.

This way, the identity of ellipse *i* as “seed” can be established based on a threshold of overlapping. This is illustrated in Fig. 8B, where ellipse *i* is selected as seed because it overlaps less that an given value with any ellipse in plane z-1. Finally, a less restrictive approach is to identify every ellipse in plane z that has not been selected as part of any 3d-object originated in the above planes. This methods is more suitable for samples with regions of high density of nuclei, and results in a larger number of “seeds” that can generate new 3d-objects. This last approach is the one used in the present contribution to quantify the digital images, as well as the zebrafish developing retinas.

In all previous approaches, all ellipses in the *z* = 1 plane are identified as “seeds”. Next, starting from the “seeds”, the framework starts to perform the reconstruction of the 3d-objects in the sample based on a combination of nonlinear fitting and statistical analysis. Identifying the correct section in the plane for each 3d-object is also important, not only to establish the number of objects, but also their location in the 3D space.

A first approach in this process is to assume that, since the ellipses fit the confocal sections of the objects, that each ellipse can be part of a single 3D-Object (illustrated in Fig. 8D). Although this a valid approach when resolution and signal-to-background ratio is high in all three spatial orientations, it is not useful in more realistic conditions when segmentation is far from perfect. To overcome this, the framework allows that each fitted ellipses can be part of several 3D-objects at the same time (illustrated in Fig. 8E). This way, the framework is able to bypass potential errors in the segmentation of the planes of the image that are typical in situations where objects are very close to each other.

Next, a key step in the process is to identify the correct sections of a given 3D-object, among all potential ellipses of a given plane. To do that, the framework computes mathematical operations using one or a combination of several parameters of the fitted ellipses to establish a ranking of all the potential interactions between ellipses in adjacent planes. Some of this potential methods to rank interactions and select the correct ellipses are defined below (and illustrated in Fig. 8).

- The simplest scenario uses the Euclidean distance between centroids to rank the interactions between an ellipse *i* in plane z-1 and the ellipses in plane z (illustrated in Fig. 8A). This method simply computes the distance between the centroids of ellipse *i* and the k-closest ellipses in plane z (up->down, left scheme). Another approach consist in selecting the k-ellipses in plane z that rank ellipse *i* as one of their k-closest in the plane *z* − 1 (down->up, center scheme). A more robust approach is to combine both (up<->down, right scheme), *i*.*e*., rank ellipses based in the distance between *i* and its *k*-closest, ans selecting only the ones that have *i* as one of their *k*-closest.
- Another approach is to rank based on the intersection area between ellipses to establish interactions (Fig. 8B). This way, we can use the percentage of area that ellipse *i* in *z*−1 intersects with each ellipse in the plane z (up->down, left scheme), or the percentage of the area of the ellipse *j* in *z* that intersects with each ellipse in *z* − 1 (down->up, center scheme), computed as explained previously. Calculation of the intersection *U*_*i,j*_ between two ellipses *i* and *j* is computed as described above when defining the “seeds”. Unfortunately, this method favors the selection of ellipses of larger size (that may overlap more with ellipses in planes above). To overcome this, the two previous methods are combined (up<->down, right scheme) to compute a more robust approach that is less biased towards larger ellipses. This way, we can define an intersection index *II*_*i,j*_ between ellipses *i* and *j* as:

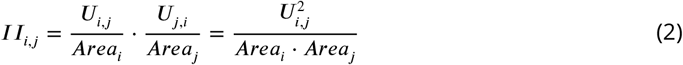

which is true because the intersection *U*_*i,j*_ = *U*_*j,i*_. Therefore, the algorithm only requires a single computation of the overlapping area.
- Although the method of areas is robust and very informative, the algorithm is not optimal in terms of computing power and memory consumption. Therefore, a faster approach that produces similar results is to rank based on the percentage of perimeter of ellipses that is inside the area of other ellipses (Fig. 8C). In this case, we compute the number of points from the perimeter of ellipse *n* fall inside ellipse *m*. Similarly to previous cases, this method can be used in (up->down, left), (down->up, center) or (up<->down, right scheme). The later is more reliable since it corrects for the relative size of the objects being compared.
- In addition, ellipses in z can be ranked based on the history of the interactions of ellipse *i* with ellipses in the previous planes. To do that, we fit the location of the centroids to a 3D line using the ‘least squares method”, i.e, combining two linear regressions over the plane *xz* and over the plane *yz*. The line that results from the intersection of the two planes represents the axis of the 3D-Object of that the ellipse *i* is part of. Finally, the ellipses in *z* are ranked based on the Euclidean distance to the point of intersection of the axis and the *z* plane, which corresponds to the expected location of the object in the *z* plane. This method is specially suited for situations where objects can be tilted with different orientations in in the 3D space, as it occurs in many biological systems, such as the zebrafish retina. An illustration of this process is shown in Fig 8D-E.
- Finally, a more complex, robust and versatile approach consists in combining several of the methods described above. This allows to filter and rank the ellipses of the plane z in sequential steps, resulting in a more reliable prediction of the interaction of ellipse *i*, that is useful for resolving 3D images of very different features. This situation is illustrated in Fig 8F, where the first k-closest neighbors are chosen based on simple Euclidean distance (such as in Fig. 8A). Next, these neighbors are tested for overlapping of perimeter (such as in Fig. 8C), and the ones that do not overlap a certain amount are discarded from the analysis. Finally, the chosen ellipse is the closest to the projection of the axis of the 3D object (such as in Fig. 8E).

The final step in the 3D-object reconstruction process is to define when a given ellipse *i* is the end point of a given 3D-Object. This is again, a key step of the process that strongly impact the number of objects that are being detected. As first approximation the tool considers an end point when a a given ellipse *i* that is part of a 3D-Object does not interact with another ellipse in the next plane (illustrated in Fig. 8A.

Although this is sufficient when working with good quality images of sparse cells, the decision of when a given section is the end point of a given 3D-Object is not so direct, since biological samples are often very dense and lack this near-perfect resolution. In this conditions, the framework uses uses statistical analysis and curve fitting for establish end points of 3D-Objects. To do that, we first estimate the average size of the objects in the *z* direction (*z*_*mean*_). We do that by estimating the minimum Feret diameter for each section of the object in all planes of the image. Next, we generate a new binary image where each section is represented as a circle with diameter equal to it minimum Feret diameter. Next, the 3D-image is processes using a smooth filter in *xz* plane. The *z*_*mean*_ value is finally established as the mean height of the objects in the *xz* plane.

The *z*_*mean*_ value is then used to estimate the end section for a given 3d-object using different approaches. The first possibility is to use the first 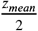 sections of a 3D-object to perform a curve fitting in three dimensions (illustrated in Fig. 8B for 3d-ellipsoids, that can be used to fit cellular nuclei). The fitting is then used to estimate where in *z* the 3D-object is expected to end, and to identify this way the last section of the object.

When images are highly distorted and signal-to-background is so reduced that results in multiple segmentation errors per object, the approach used is based on the characterization of the integrity of each 3D-object constructed. This integrity is estimated statistically based on three parameters: 1) the euclidean distance of each centroid of the 3d-object to the 3D projection line (as defined above) in each plane; 2) the angle between the 3d-projection line and the vectors formed by two consecutive points, 3) the cumulative Pearson Correlation Coefficient *ρ* between the two previous variables.

In addition, in sharp objects that can been imaged at high resolution and signal-to-background, both distance and angle should remain close to zero, and *ρ* ;≈ 1 (the two variables are perfectly linearly correlated). On the other hand, angle and distance are higher in poorly segmented objects, and they become uncorrelated (*ρ* → 0) as we sequentially add more sections to the 3d-Object).

Using these values, we define the end point of a 3d-Object as the last centroid in a 3d-object that remains inside a 50% confidence interval for all the three parameters. The 50% confidence interval is calculated from the distances and angles of the first 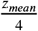 planes.

Based on this value, for an 3D-object composed on *n* planes, we can compute the number centroids inside 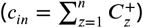 or outside 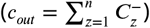 the value of CI.

Finally, we establish the end of the 3D-object in the last centroid that fulfill the following inequality.

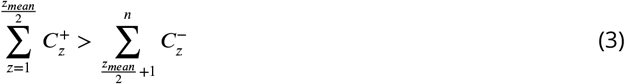

For sharp 3D-objects, the end point corresponds to the last centroid that lies inside the confidence interval (illustrated as the cylinder in 8C), while in noisy objects, some centroids in the first 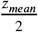 sections can be outside the confidence interval, so the cut occurs when more centroids after *z*_*mean*_/2 are are outside CI than in the first *z*_*mean*_/2 planes (illustrated in 8D). This is computed as a filter that allows us to estimate the end section of a 3D-object in conditions of low signal-to-background.

Finally, if the integrity of the 3D-object is so compromised that the previous equation is not fulfilled even after reaching a number of planes equal to 2 × *z*_*mean*_, then we assume that we are unable to find the correct end point of the present 3d-object. In this case, the 3d-Object is assumed to end at the plane *z*_*mean*_.

### Module 4: Establishing nuclei identity based on immunostaining

The high accuracy in object detection of OSCAR is used to extend its capabilities of and implement accurate automated fate identification of the cells in the three-dimensional space. Cell identity is established based on the intensity of the immunofluorescence staining against markers characteristic for each particular fate. To perform this task, the samples are stained *in toto* following the protocol described above, cleared and mounted. Next, during the image acquisition stage, a multi-dimensional image is obtained with multiple laser wavelengths simultaneously, so each color channel corresponds to a different immunostaining of the sample.

Next, each color channel of the image is processed independently to enhance contrast and reduce noise. An intensity value is defined based on the particular features of each immunostaining for to establish the threshold between signal versus background.

Next, for each individual 3D-object in the image, the total volume is computed as the sum of the areas *A*_*i*_ of the ellipses of its sections.

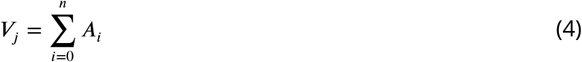

where *i* = 1…*n* indexes all the ellipsoidal sections of each 3D-object.

Next, we establish a threshold value for each channel, to distinguish clear positive staining with background levels. Next, we compute the number of pixels in all ellipses 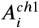 of the object that are above the threshold for each channels. The sum of all these areas is defined as the volume 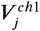

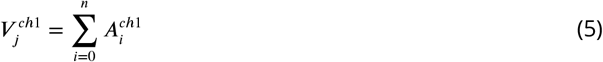

Finally, object *j* is identified as positive or negative for each particular staining if 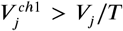, where the free parameter *T* is adjusted depending on the features of the image and the staining. For the experiments analyzed in this manuscript, the value is set to *T* = 4 (at least one out of 4 pixels for object *j* is above the threshold for this particular staining).

When the identity of object *j* can be established based on two complementary fluorescent labeling (such as in the combination between SOX2 and GPF to establish identity as progenitor or differentiated cell), an additional layer of robustness is added by comparing the amount of pixels positive of each channel, so if 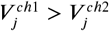, then the object is considered positive for immunostaining of channel 1, while if 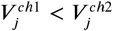, the object is considered positive for immunostaining of channel 2. For the object with similar number of pixels above the intensity threshold (i.e., 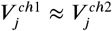), the decision is made based on additional features (such as in this contribution, by measuring staining against Phospho histone 3 and PCNA).

### Generation of output figure

Final 3D representation is done using the tools of Fiji 3*Dviewer* and 3*D* − *DrawShape* from the package 3D ImageJ Suite (***Schmid et al., 2010***; ***Ollion et al., 2013***). Lookup Table (LUT) for each image is modified at convenience (color legend is indicated in each figure). Output image is generated by drawing objects of a given shape in the location of the centroids of each 3*D* − *object* calculated. In this contribution, nuclei are represented in the output image as ellipsoids where dimensions are obtained from the parameter of the 3*D* − *object* it represents: *x* − *length, y* − *length, z* − *length*, and orientation (represented for the 3D-Object line). Objects in the output image are represented with different colors based on the nuclei identity calculated in the previous section.

### Accuracy detection in the 3D space

To statistically asses how good is the spatial location of each tool, the Friedmann-Rafsky (FR) test is used. The FR test is an statistical method to tests whether two point clouds belong to the same distribution. The test is a generalization of the Wald-Wolfowitz runs test for *n >* 1 dimensions and its based in computing the Minimum-Spanning Tree (MST). In brief, the tool generates a connected, edge-weighted and undirected graph from two sets of points distributed in the 3D space, in which each pair of nodes is connected in such way that all the vertices remain connected but without any cycle. The graph is constructed in such a way that minimizes the total Euclidean distance (considered as the weight of the edges connecting the nodes).

Then, the number of “runs” (*R*) is computed as the number of edges connecting different samples plus one. Finally, the FR-statistic *W* is calculated as:

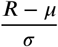

being *µ* and *σ* the mean and standard deviation of the *R* distribution (***Chen et al., 2018***; ***Friedman and Rafsky, 1979*)**.

Since the result depend on the size of the samples, to reject the null hypothesis, with bigger samples the rejection *p* − *value* becomes smaller.

On the other hand, randomization of the subsample forces to run the test several times and compute a median to avoid biases. Common approaches consist in taking random subsamples of the same size for both samples and running the test as many times as points taken, then, the rejection *p* − *value* is adjusted.

As we are not interested in rejecting the null hypothesis but in comparing normalized FR statistics for the different methods, we have chosen to compare 500 random centroids of the objects found against the whole set of centroids of the artificial image. As *W* follows a standard normal distribution, the lower the statistic the better, i.e. smaller values mean higher degree of similarity between distributions (***Hsiao et al., 2016***) and, thus, better spatial location of the found objects.

### Software used and comparison with OSCAR

#### Object Counter 3D

(OC3D) is an open-source algorithm implemented in Fiji ((***Bolte and Cordelières, 2006*)**, https://imagej.net/3D_Objects_Counter). The principle of OC3D to localize objects in 3D images is based on voxel connectivity. In other words, OC3D identify as single 3D object all voxels with an intensity value higher than a certain cutoff value (foreground) that are connected without interruption of voxels that do not are higher than the cutoff value for voxel intensities (background). The detection of the background implies the end of a detected 3D object.

#### TANGO

The plugin “3D analysis” integrated in the package 3D ImageJ Suite of Fiji (TANGO) is highly similar in terms of accuracy to OC3D, however TANGO employs a different way to calculate spatial and morphological measurements of each 3D objects (including the centroid) ((***Ollion et al., 2013***), https://imagej.net/3D_ImageJ_Suite)). TANGO works only with images whose 3D objects have been identified and labelled previously. Each label is a specific voxel intensity associated with only one 3D object, therefore TANGO only needs to look in the histogram of voxel intensities to identify them.

Both TANGO and OC3D in this comparison use as input images that have been segmented using the algorithm 3D-watersheed from the package 3D ImageJ Suite (accuracy for images using previous 2D-watersheed or without any watersheed is always lower than the 7%).

#### Imaris© 8

(BitPlane, https://imaris.oxinst.com/) is a commercial platform of image visualization, processing and analysis, widely used by the scientific community and with an easy interface that requires no programming skills and presents a large variety of tools. Data from Imaris© is used after activating for each tool the segmentation step and the 3D detection tool available, often used in other other studies ((***Matejčić et al., 2018***)).

#### Huygens©

Huygens Professional version 19.04 (Scientific Volume Imaging, The Netherlands, http://svi.nl) is another commercial platform,

### Branching Process tool

Average cell cycle length and mode of division are measured based in a tool developed by our lab (***Míguez, 2015*)**. The tools is composed of simple analytical equations derived using a branching process formalism. Input of the equations are the number of progenitors and differentiated cells overtime, the growth fraction and the apoptosis rate. The equations are :

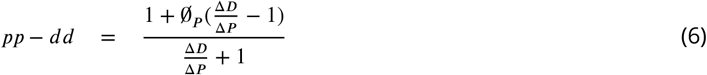

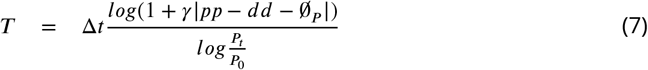

The value *pp* − *dd* can be identified with a measure of the differentiation dynamics (*pp* and *dd* are the rate of symmetric proliferative and differentiative divisions, respectively), and goes from 1 (all divisions being symmetric proliferative) to -1 (all divisions being symmetric differentiative). This way, *pp* − *dd* = 0 corresponds to progenitor pool maintenance, which can be achieved via asymmetric *pd* divisions (if *pd* = 1, then *pp* = *dd* = 0) or via balance between symmetric proliferative and differentiative divisions (*pp* = *dd* independently of the value of *pd*). Unfortunately, by construction, the model does not distinguishes between these two mathematically equivalent scenarios).

Δ*P* = *P*_*t*_ − *P*_0_ and Δ*D* = *D*_*t*_ − *D*_0_ are the number of progenitors and differentiated cells generated in a given window of time Δ*t* = *t* − *t*_0_. ∅_*P*_ corresponds to the rate of cell death of the progenitors. *γ* is the growth fraction (*γ* = 1 if all progenitors are actively cycling).

## Notes

### Competing Interest Statement

The authors have declared no competing interest.

